# Adaptive gene duplication can exceed optimal dosage

**DOI:** 10.64898/2026.09.29.755440

**Authors:** Frédéric Bédard, Alexandre K. Dubé, Isabelle Gagnon-Arsenault, Christian R. Landry

## Abstract

Gene duplication is an important evolutionary mechanism contributing to genomic innovation and adaptation. While the long term fate of duplicated genes has been extensively studied, immediate effects such as those resulting from increase in gene dosage remain understudied. Of particular interest is the question of how expression level relates to the optimum after a duplication, as it could fall short, exceed or directly reach the optimal expression. In this study, we investigated the relationship between expression and fitness for 11 genes previously reported to increase fitness when duplicated in *Saccharomyces cerevisiae* to compare the expression level achieved by duplication to the one maximizing fitness. While most of the studied genes show little fitness variation across the expression levels tested, we found that *ESP1*, which encodes a separase essential to anaphase initiation, exhibits a clear fitness optimum at an expression level above the native single-copy state. However, the expression level associated with gene duplication exceeds the optimum. We find that *ESP1* overexpression is associated with a reduced cell size and a lower proportion of the cell cycle spent in G2 phase, consistent with its role in cell cycle progression. We propose that the duplication of *ESP1* accelerates the cell cycle, resulting in smaller, faster-dividing cells. When measured in different environments, the fitness functions also exhibit similar shapes, with slight variation in the optimal expression. Our findings highlight how gene duplication can confer an immediate selective advantage through dosage increase while being non-optimal, reversing the directionality of natural selection on expression level after duplication.

## Introduction

Gene duplication is a major mechanism for the emergence of new genes and thus potentially new biochemical functions. Throughout the past decades, different evolutionary mechanisms that explain the retention of paralogs have been examined in detail. These include the emergence of new functions (Ohno, 1970), the escape from adaptive conflicts (Des Marais & Rausher, 2008), sub-functionalization (Force et al., 1999) and paralog interference (Cisneros, Mattenberger, et al., 2026; Kaltenegger & Ober, 12/2015). While all these mechanisms lead to the long-term retention of paralogs, the functional divergence required for these mechanisms to take place may require time and cannot occur if the duplication is rapidly lost in the population (Lynch & Conery, 2000). Immediate effects that would render gene duplication beneficial therefore play a crucial role in the evolutionary fate of new duplicate genes, since they would increase the probability of fixation and their following maintenance (Kondrashov & Kondrashov, 2006).

One immediate consequence of gene duplication relates to expression. Following its duplication, the expression level of a gene may increase through change in gene dosage (Gu et al., 2005; Loehlin & Carroll, 2016; Rogers et al., 2017). While this can have deleterious effects for certain genes or be neutral, increased dosage can also result in a fitness advantage immediately after duplication (Kondrashov & Kondrashov, 2006) (Fig. 1A). Under the “positive dosage” model of gene duplication, increased expression has been suggested to be beneficial for genes involved in metabolism (Perry et al., 2007) stress responses (Hardwick et al., 2014; Robinson et al., 2023; Selmecki et al., 2008), and for genes for which the product is necessary in large amounts under certain conditions (Kondrashov, 2012). For example, increased copy number of the salivary amylase gene (*AMY1*) is observed in human populations with starch-rich diets (Perry et al., 2007). In microbes, increased gene copy-number can cause drug resistance by increasing the dosage of a drug target (Todd & Selmecki, 2020).

**Figure 1.**
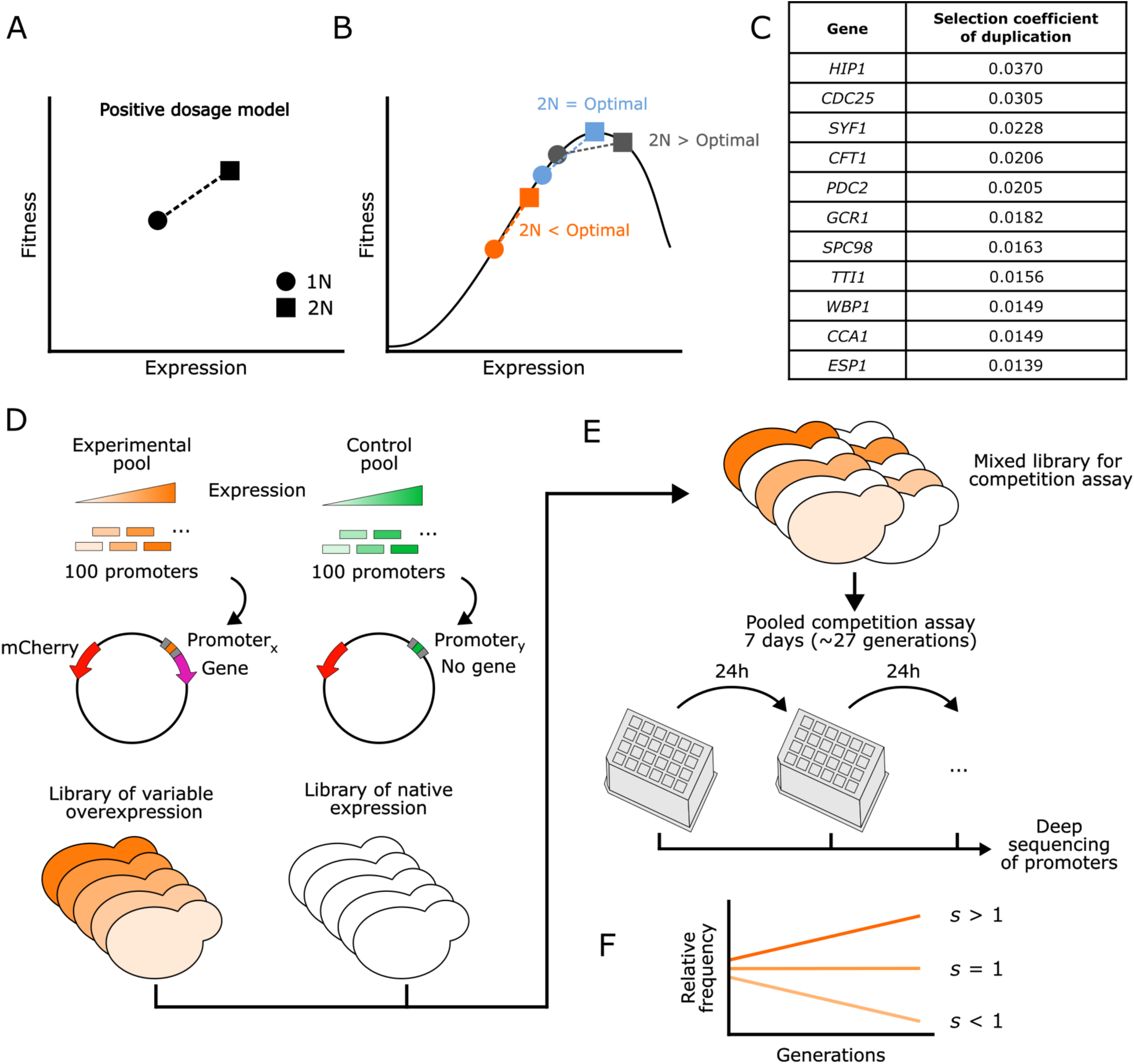
Experimental system to study fitness functions above native expression levels. A) Schematization of the positive dosage model. Gene duplication (2N) can result in higher fitness than the native state (1N) through an increase in the gene product concentration. B) Increased fitness resulting from gene expression increase could result from different expression levels compared to the optimal level, either by reaching it directly (blue), or getting closer to the optimum either by not reaching it (orange) or exceeding it (black). C) Eleven yeast genes presenting a positive selection coefficient when duplicated were selected from Ascencio et al. (2021). D) For each gene, a library of 100 plasmids carrying a gene copy controlled by synthetic promoters of varying strength was transformed in the R1158 *S. cerevisiae* strain, resulting in a library of genotypes with different expression levels in addition to the endogenous gene copy. A library of 100 plasmids with different but equivalent synthetic promoters, but with no additional gene copy, was also transformed in the same yeast background as a control library. E) For each gene, the experimental and control libraries were mixed and pooled competition assays were performed. Deep sequencing of the promoters at each timepoint allowed the tracking of each genotype’s frequency through time. F) The selection coefficient (*s*) of a genotype is calculated from its change of frequency relative to the controls throughout the experiment.

While immediate beneficial effects have been observed following the duplication of several genes (Ascencio et al., 2021), little is known regarding the extent to which gene expression is increased, and what mechanism results in the observed increased fitness, i.e. what cellular phenotypes lead to faster growth rate. A fitness increase suggests that expression level was originally suboptimal. Although this implies that duplication brings the expression of a gene closer to its optimum, whether the exact optimal level is reached or whether further increase or decrease of expression would lead to an even higher fitness remains unclear (Cisneros, Dibyachintan, et al., 2026). This can be illustrated with a fitness function, which relates fitness to gene expression (Fig. 1B). Accordingly, a better understanding of the role that gene expression increase can play in immediately beneficial duplications requires investigating these fitness functions. If the expression level remains suboptimal, natural selection should favor the evolution of expression to reach the optimum over time. This optimization of gene expression has notably been shown by the rapid evolution of the expression of the *lac* operon in *Escherichia coli* to match the optimal protein abundance (Dekel & Alon, 2005). This implies that the evolutionary trajectory of dosage following gene duplication could be impacted by the shape of the fitness function, the different scenarios presented in Fig. 1B resulting in different selective pressure on gene expression after gene duplication. Natural selection would favor an increase in expression if duplication led to sub-optimal expression levels while it would favor a decrease if the optimum was overshoot.

Synthetic biology tools now enable the modulation of gene expression to different levels and the measurement of their effect on fitness, allowing the investigation of fitness functions and the positioning of gene duplication on these functions (Arita et al., 2021; Keren et al., 2016; Otto et al., 2024; Sopko et al., 2006). In this study, we set out to examine if beneficial gene duplication leads directly to optimal expression or if it overshoots or falls short of this optimum. To do so, we examined the fitness function of 11 genes for which duplication was reported to provide a selective advantage upon duplication in a large-screen performed in *Saccharomyces cerevisiae* (Ascencio et al., 2021) (Fig. 1C, Supplementary Table 1). Using a library of 100 synthetic promoters of graded strength, we individually modified the expression of each gene and established their fitness functions with pooled competition assays (Fig. 1D-F). This allowed us to compare the optimal gene expression with the level reached following gene duplication. While most genes show little effect of expression on fitness in the range of expression levels tested, we found that for *ESP1*, a gene involved in cell cycle regulation, duplication leads to an expression level exceeding the optimum. The fitness function of *ESP1* is a concave parabola, with a positive correlation between fitness and expression below the optimum and then a negative correlation above. The phenotypic analysis of cells showed that increased *ESP1* expression is associated with smaller cell size and a reduction of the ratio between the duration of the G2 and G1 cell cycle phases. This led us to propose that the selective advantage associated with the duplication of *ESP1* could be the result of an acceleration of the cell cycle, providing a mechanistic explanation of how gene duplication can lead to an immediate selective advantage through increased expression.

## Results

### Expression modulation using synthetic promoters

We studied the fitness function of 11 genes identified in Ascencio et al. (2021) as leading to higher fitness when duplicated on a centromeric plasmid. These fitness effects were determined by conducting competition assays, which compared, through time, the frequency of a strain carrying an additional gene copy on a plasmid with the one of a control strain carrying an empty plasmid. Each strain was marked with a different fluorescent reporter allowing the measurement of the change in the proportion of the two genotypes through time. The selection coefficients associated with the duplication of the selected genes ranged from 0.0139 to 0.0370 (Fig. 1C). These genes cover different molecular functions (e.g. amino-acid transporter, cell division, transcription factors) and a range of protein abundance (Supplementary Table 1).

To establish the fitness function of these genes, we used an approach in which a strain of *S. cerevisiae* carries an additional copy of the gene of interest on a centromeric plasmid controlled by a library of 100 synthetic promoters driving a range of expression levels (referred as “pSyn1” to “pSyn100” in rank order of expression strength) taken from de Boer et al. (2020). These synthetic promoters are composed of a scaffold of 143 basepairs (bp) (referred to as “pTpA”) in which a sequence of 80 bp is cloned to form the promoter. This design enabled us to obtain 100 genotypes with varying expression levels above the endogenous gene copy for each gene of interest, as the expression level results from the sum of the genomic and plasmid gene copies. Because the promoter sequences are short, unique sequences, their frequency can be followed by deep sequencing. This allowed us to measure fitness for each genotype as they compete in the same culture, and, from these values, obtain the corresponding fitness functions.

To validate the range of activity covered by the synthetic promoters, we selected a subset of 11 promoters with activity levels distributed across the range of expression previously reported and measured their corresponding activity using a yeGFP reporter and flow cytometry. Comparing the expression measurements to the estimates of de Boer et al. (2020), we observed a perfect conservation of rank order across the measured strength of the promoters (Spearman correlation, *r*(9) = 1.00, p < 0.001) with a slight deviation from linearity for the strongest promoter (Fig. 2A). We also tested if the range of activity of the synthetic promoters covers the activity levels of the native promoters of the studied genes (1000 bp upstream of the coding sequence of genes). We measured the activity of these wild-type yeast promoters by cloning them on the same plasmid upstream of the yeGFP. Comparing their activity levels with the ones of the lowest (pSyn2) and strongest (pSyn100) synthetic promoters enabled us to confirm that the set of synthetic promoters should cover the activity levels of the endogenous promoters of the 11 genes (Fig. 2B). These results suggest that adding a plasmid copy of the genes of interest controlled by these synthetic promoters should cover a vast range of expression levels, including the ones expectedly reached by duplication and slightly beyond.

**Figure 2.**
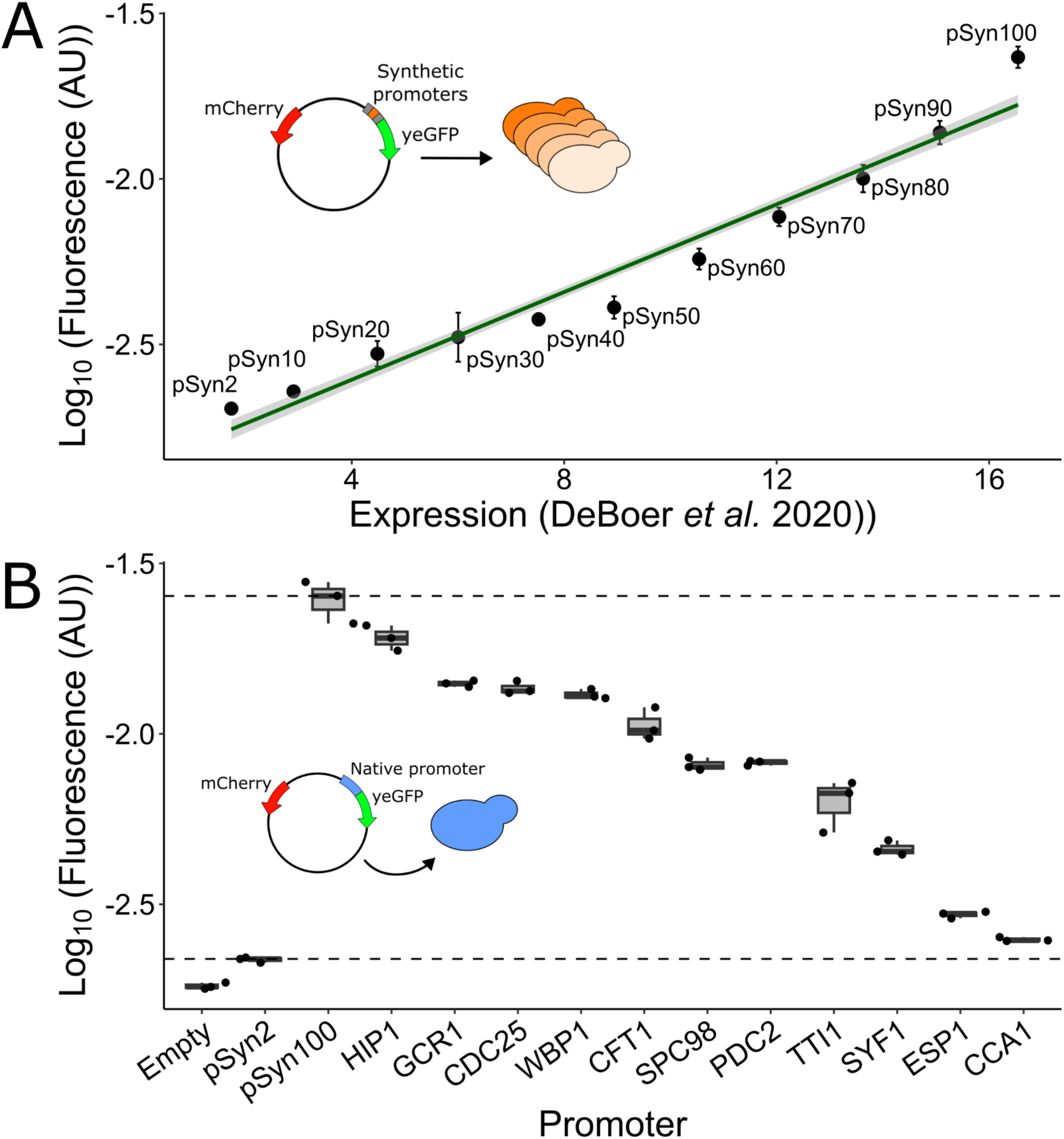
A pool of 100 synthetic promoters covers the range of the native promoter activity of the genes of interest. A) Flow cytometry measurements (mean ± standard deviation, n = 8) of the fluorescence of cells expressing yeGFP (corrected for cell size (FSC)) from 11 synthetic promoters (y-axis) distributed across the previously reported expression estimates of de Boer et al. (2020) (x-axis). Synthetic promoters are referred as “pSyn”, the following number indicating its expression estimate rank in ascending order across the pool of 100 promoters selected. B) Measurements (n = 3) of the activity of each gene’s native promoter upstream (1 kb upstream of the start of the coding sequence) of yeGFP on the same plasmid as in A) through flow cytometry fluorescence measurements (corrected for cell size). The median strength of one of the lowest and strongest promoters (pSyn2 and pSyn100 respectively) are represented as dashed lines and indicate the limits of the range of detectable expression of the synthetic promoters.

### Fitness assessments of strains expressing an additional copy of the genes

To estimate the fitness function of each gene beyond the single-copy expression level, we performed competition assays with a strain transformed with the pool of 100 plasmids expressing the gene controlled by the set of promoters. As controls, the same strain background transformed with another pool of 100 plasmids with synthetic promoters also covering the same range of activity, but without an additional gene copy, was pooled with the ones carrying the gene inserts. These offer a reference set of plasmids with various promoters but no additional gene expression, reflecting the fitness of the single gene copy that is in the genome. The assays were performed in synthetic media for 7 days (∼27 generations) without agitation, diluting the cultures by a factor of 1:25 in fresh liquid media every 24 hours. The assay was performed for each gene in separate cultures in which promoter genotypes are competing with each other. These conditions were chosen to replicate the conditions of Ascencio et al. (2021) as much as possible. Before each dilution throughout the assay, we sampled the culture and measured the frequency of each promoter genotype using deep sequencing. This allowed the calculation of a selection coefficient associated with each promoter (methods). This assay was separately conducted for each gene in triplicate, allowing the association of selection coefficients to the relative expression levels provided by de Boer et al. (2020). This enabled us to establish the fitness function over the expression range added from the plasmid gene copy (Fig. 3), which was drawn by fitting a LOESS regression.

**Figure 3.**
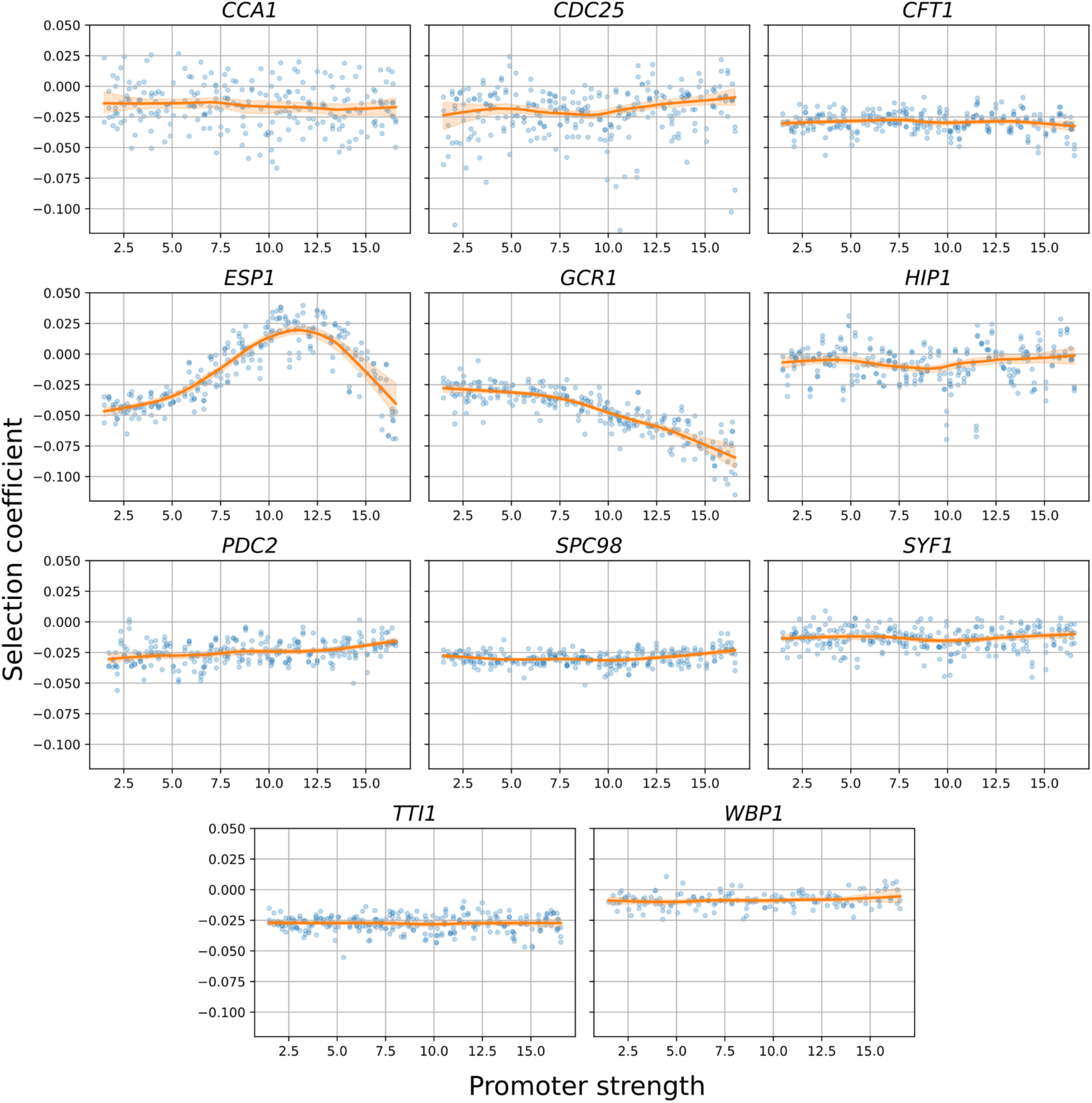
Fitness functions of the 11 genes of interest. The relationship between fitness and the expression levels driven by the synthetic promoters. The x-axis corresponds to the gene expression level reported in de Boer et al. (2020) from the plasmid copy, which is additional to the endogenous nuclear gene copy. Fitness functions were obtained using a LOESS regression (frac = 0.4) and shown in orange, whereas the shaded areas represent the bootstrap 95% confidence interval (n = 200). Selection coefficients were calculated for each replicate (n = 3) of each promoter and are shown in blue. Correlation between replicates for each promoter and selection coefficients for the control pool are available in the Supplementary Material (Fig. S2 and S3 respectively).

While the fitness functions of most genes appear flat in this range of expression, the fitness function of *ESP1* presents a clear fitness optimum, from which increasing or decreasing expression results in a fitness decrease. We also observed a negative relationship between the selection coefficient and expression for *GCR1*, going against our expectation of increased fitness with overexpression. To verify the presence of more subtle fitness effects through the promoter range across the other genes, the selection coefficient values of the five promoters with the lowest and highest strength were compared. This comparison was not significant for all genes except *PDC2,* for which the selection coefficients of the strong promoters are significantly higher than the ones of the weak promoters (t-test, *t*(8) =-3.03, p = 0.016), which indicates the presence of a fitness increase with overexpression. As no plateau could be identified for the fitness functions of this gene, the optimal expression level was not observed, suggesting that optimal expression would be higher than the range explored with the set of promoters.

We also noted that the fitness functions present an initial negative selection coefficient for the weakest promoter as compared to the control set of plasmid (not containing any gene inserts). As this is unlikely to be caused by the weak expression of the plasmid gene copy, we suspected a bias between the control and experimental plasmids. Since the sole difference between the two types of plasmids is the presence or absence of the gene of interest, we hypothesized that this bias could be linked to gene length, as plasmid size has been reported to affect the length of the lag phase in *E. coli* (Cheah et al., 1987; Smith & Bidochka, 1998). Comparing the initial fitness of the fitness function of each gene, we indeed found that the selection coefficients of the weakest promoter significantly correlates with the length of the gene added on the plasmid (Spearman correlation, *r*(9) =-0.691, p = 0.019) (Fig. Sup. 1), suggesting that increased plasmid size through the addition of the gene results in a proliferation disadvantage for the experimental genotypes. While this prevents us from comparing the selection coefficients of the experimental pool to the ones of the controls and to consider the latter as the single copy state, this bias does not affect the shape of the functions, since all experimental genotypes carry a plasmid of the same length.

Although our set of selected genes were reported to exhibit fitness advantage when duplicated (Ascencio et al., 2021), most of the fitness functions exhibit little variation with expression. We therefore hypothesized that while the native promoters have activities falling within the range of the synthetic promoters, protein abundance resulting from the latter may not reach the level attained when the genes are duplicated with their native promoters. We tested this hypothesis by tagging the genomic copy with yeGFP and constructing an empty plasmid (no gene copy or yeGFP), as well as plasmids with the gene copy tagged with yeGFP and controlled either with its native promoter or the second most potent synthetic promoter (pSyn99). When transformed in the yeGFP tagged strain, these three constructs respectively allowed the measurement of expression of the native gene, the native gene when duplicated, and the maximal expression reached with the synthetic pool. Through flow cytometry measurements (Fig. 4), we observe that the maximal protein abundance reached for 6/10 tested genes is higher than the one at gene duplication. As only the promoter and gene coding sequence were duplicated, the absence of the terminator region, present in the duplication constructs of Ascencio et al. (2021), might influence the protein abundance measures of the duplicated state, some regulation mechanisms potentially absent in our setting. Nonetheless, in the case of the four genes for which the protein abundance at the duplicated state was not reached with the strongest synthetic promoter (*CDC25*, *GCR1*, *HIP1* and *WBP1*), different regulation mechanisms probably reduce the protein abundance below the duplicated state, even though the synthetic promoters might bring the transcription level of our set of genes above the one of the duplication. For these genes, the tested range of protein abundance would therefore be too low to measure potential beneficial effects at the duplicated state, instead resulting in flat fitness functions. Contrastingly, the slight fitness increase with the overexpression of *PDC2* without reaching a fitness plateau and the confirmation that the range of expression covers the one reached with its duplication suggests that the optimal gene expression would be beyond the duplicated state.

**Figure 4.**
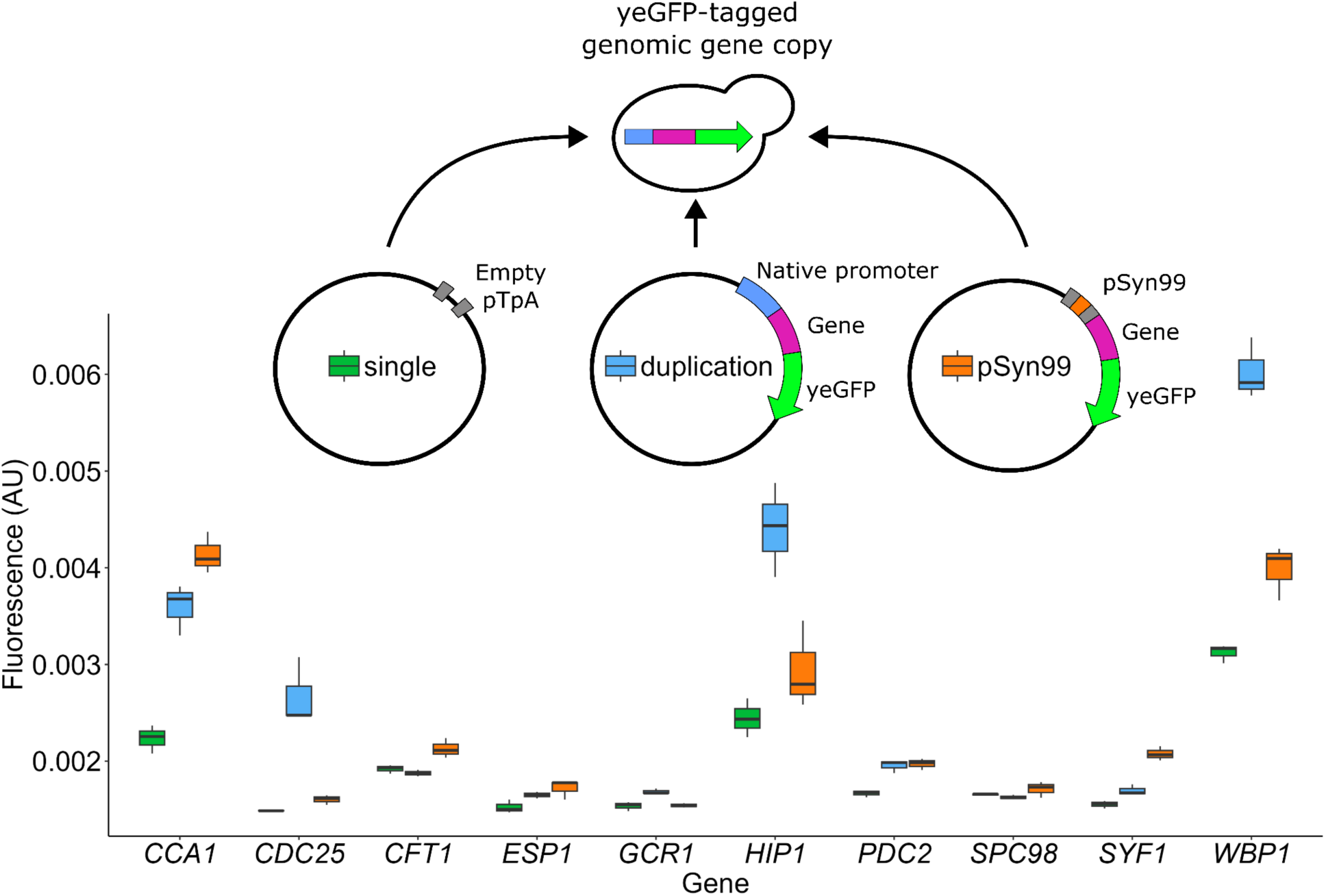
Measurements of total expression of various constructs to compare protein abundance driven by native and synthetic promoters. Single copy (genomic copy fused to yeGFP and empty plasmid), duplicated (genomic copy fused to yeGFP and plasmid copy with endogenous promoter and coding sequence fused to yeGFP) and overexpressed with pSyn99 (genomic copy and plasmid copy with one of the strongest synthetic promoters driving the expression of the coding sequence fused to yeGFP) states for 10 of the studies genes. Measurements were done in triplicate by flow cytometry, and yeGFP fluorescence was corrected for cell size (FSC).

### The duplication of ESP1 overshoots the optimal gene expression

Since the fitness function of *ESP1* shows a clear optimal expression level within the covered range of overexpression, we focused on this gene to examine if duplication results in an expression level close to the optimum. We tagged both genomic and plasmid gene copies with the yeGFP and measured Esp1p abundance with either the native promoter or different synthetic promoters on the plasmid, using pSyn65 as the optimal expression level (Fig. 5A). Our results show that the expression level reached through duplication is slightly higher than the optimum (t-test, *t*(14) =-2.15, p = 0.049) (Fig. 5B). This suggests that for *ESP1*, the fitness advantage of gene duplication occurs even though it slightly overshoots the optimal expression level.

**Figure 5.**
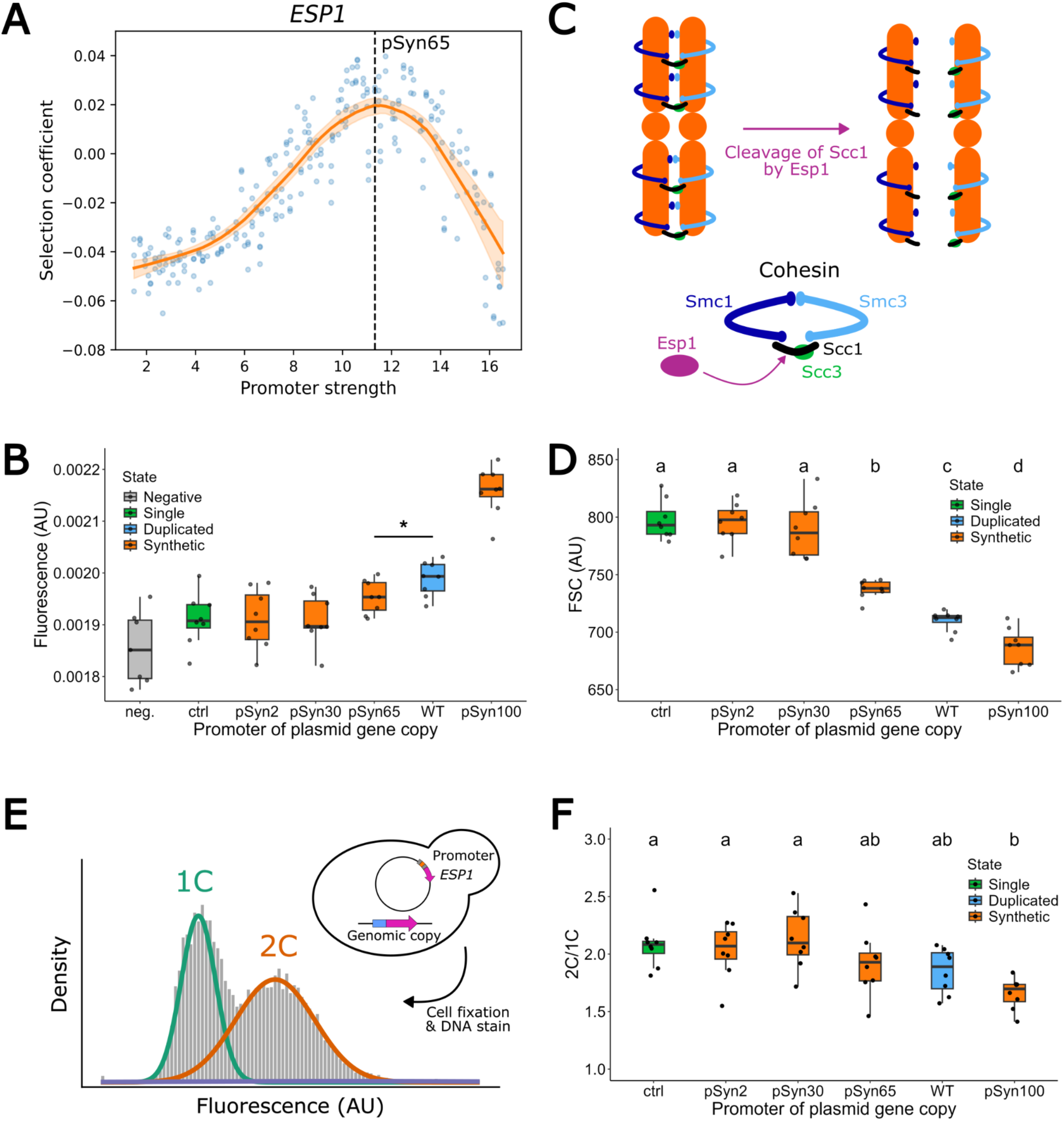
The duplication of *ESP1* overshoots the optimal expression levels and is associated with reduced cell size and G2/G1 proportion in the cell cycle. A) pSyn65, identified as the dashed line on the fitness function of *ESP1* produces optimal expression level for *ESP1*. B) Esp1p expression measurements (n = 8) associated with gene duplication (genomic and plasmid gene copy controlled by the endogenous promoter) and overexpression levels reached with different synthetic promoters. Both genomic and plasmid copies were fused to yeGFP to measure expression through fluorescence corrected by cell size (FSC). Both an untagged genomic strain (neg.) and a tagged strain (ctrl.) with a control plasmid were added, respectively representing a negative control and the native expression level. Significance of the comparison of expression levels for the promoters pSyn65 and WT is indicated as follow: (*: p < 0.05). C) Schematics of the function of Esp1p. *ESP1* encodes a separase which cleaves cohesin, resulting in the separation of sister chromatids and the initiation of the anaphase D) Forward scatter signal as a proxy for cell size measurements (n = 8) for each Esp1p expression level tested in C). Strains and plasmids were the same, with the exception of the yeGFP fusion absent for these measurements. One *ESP1* plasmid with an empty pTpA and stop codons in the coding sequence (ctrl.) was included as a control. Ratios were compared with a two-way ANOVA and a Tukey HSD post-hoc test (α = 0.05), for which the results are indicated using a letter code. E) Experimental design for DNA content measurements. The proportion of time spent in G2/G1 was measured through DNA content measurements (n = 8) of fixed exponential phase cell cultures of different genotypes. Cells were assigned to 1C (normal DNA content) or 2C (replicated DNA) with a Gaussian Mixture Model based on DNA content. F) The G2/G1 ratio decreases with the overexpression of Esp1p. Tested levels are the same as D) and statistical results are presented in the same manner.

*ESP1* encodes a separase that dissociates sister chromatids during cell division (Fig. 5C). Since Esp1p is involved in the cell cycle (Baum et al., 1988; Uhlmann et al., 1999), we hypothesized that increased expression could be associated with an acceleration in the cell cycle, resulting in faster-dividing cells but that could, as a result, be smaller. Cell size measurements using flow cytometry supported this hypothesis, as cell size decreased significantly in strains with higher *ESP1* expression levels (Tukey HSD, p < 0.05) (Fig. 5D). Given its role in the cell cycle and more specifically in the initiation of anaphase, we tested if the overexpression of Esp1p could result in a faster transition from G2 to G1. We therefore measured the distribution of DNA content in exponential phase cell cultures carrying plasmids with different promoters to test whether increased expression affects the G2/G1 ratio (Fig. 5E), the reasoning being that a change in the ratio between diploid and haploid cells would indicate a change in the proportion of the cell cycle spent in each of the two states. Using a gaussian mixture model to assign observations to either C1 (haploid DNA content) or C2 (doubled DNA content), we observed a significant decrease in the proportion of cells in C2 between the strongest and lowest expression levels (Tukey HSD, p < 0.001) (Fig. 5F). Together, these results strongly suggest that the overexpression of *ESP1* is associated with a reduction in cell size and proportion of time in G2 during the cell cycle. This provides a potential mechanism to better understand the immediate fitness advantage associated with the duplication of *ESP1* through increased dosage in these conditions, and why the highest overexpression levels could be deleterious.

While we observed a fitness increase with *ESP1* overexpression, this duplication has, to our knowledge, not been reported in *S. cerevisiae* strains or related species. However, one could argue that given this fitness advantage, gene duplication should have already occurred and increased in frequency in the past. On the other hand, the duplication of *ESP1* could be beneficial specifically for the tested conditions, but deleterious in others, since different conditions could lead to different fitness functions (Siddiq et al., 2024). To test whether growth conditions would indeed impact *ESP1*’s fitness function, we performed the pooled experiment in 6 other conditions (Fig. 6A), changing the carbon source (galactose or raffinose instead of glucose) or its concentration (1:2 dilution of the original SC media in H_2_O), adding agitation to our cultures, increasing DNA damage stress through the addition of methyl methanesulfonate (MMS) (0.001%) or the disruption of microtubule functions, and thus potentially anaphase, using benomyl (5 μg/mL). Dimethyl sulfoxide (DMSO) (the solvent used for MMS and benomyl) was also included as a control condition. This was performed for *GCR1* as well (Fig. 6B), since it also showed a strong negative response to overexpression in the initial glucose condition (Fig. 3). Since a bias between the competition and control empty plasmids was still observed to different scales depending on the condition, a baseline correction was applied to the fitness functions so that the initial selection coefficient was 0 across all conditions, allowing comparisons between them (Fig. S4). Fitness functions were also calculated for all replicates individually to allow statistical comparisons of the optimal gene expression and highest selection coefficient for *ESP1* (Fig. 6C). The uncorrected functions for all replicates are available as supplementary material (Fig. S5).

**Figure 6.**
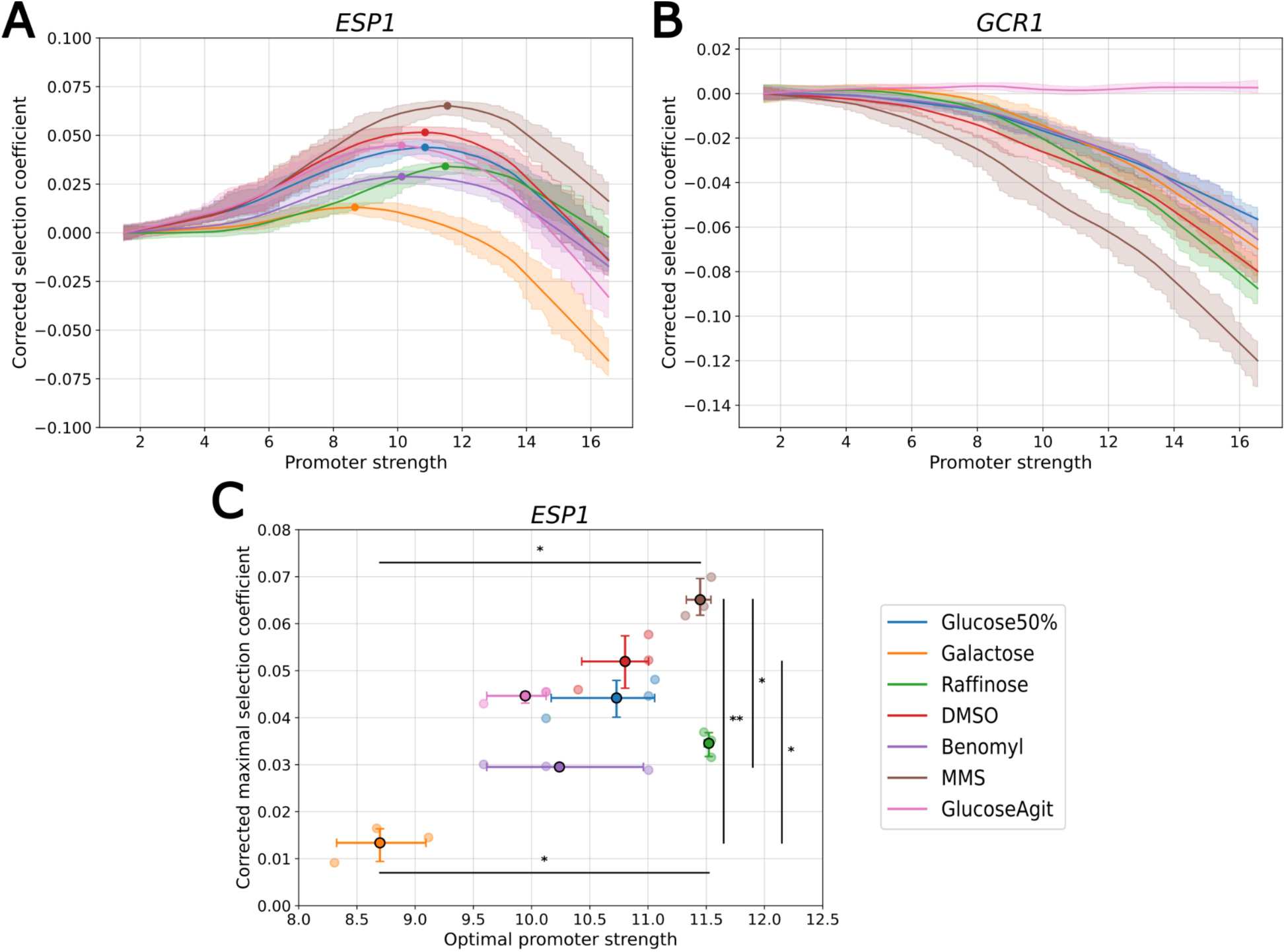
Fitness functions vary across conditions for. A) *ESP1* and B) *GCR1*. Assays were performed in triplicates for each condition (see legend, shared for panels A, B and C). Correlation between replicates for each promoter is available in the Supplementary Figure 6. Fitness functions (solid lines) were smoothed using LOESS (frac = 0.4) regression on all datapoints, with shaded areas representing the bootstrap 95% confidence intervals (n = 200). Fitness functions were adjusted to start at 0 to correct for the initial bias between the experimental and control plasmids (Fig. S4) C) Optimal promoter strength and maximal selection coefficient for *ESP1* across different conditions. Fitness functions, the optimal promoter strength and the associated selection coefficients were calculated for each triplicate of each condition (shown in faded color), with solid points representing the means. 95% confidence intervals are also provided, with significant contrasts (Dunn’s test with Benjamini-Hochberg’s correction) between optimal promoter strength (horizontal lines) and corrected maximal selection coefficients (vertical lines) identified as follows: (*: p < 0.05, **: p < 0.01).

While the shape of the fitness function remains similar for most conditions for both *GCR1* and *ESP1*, the various conditions tested alter the amplitude of the effects. For *GCR1*, we observe the general trend of negative effects of overexpression across all but one condition, the fitness function being flat when agitation is added to the initial screen conditions. This indicates that the negative relationship observed between expression and fitness for *GCR1* results from conditions related to the absence of agitation, and this effect is amplified by some of the modifications made to the media. For *ESP1*, a fitness advantage is observed across all tested conditions, showing that accelerating cell division may be generally adaptive at least for competitive fitness in laboratory settings. However, we observed significantly different optimal expression and maximal fitness advantage between some of the conditions (Krustal-Wallis test) (Fig. 6C). Pairwise comparisons using Dunn’s test with Benjamini-Hochberg’s correction indicates that optimal promoter strength was significantly lower in galactose media than in raffinose (p = 0.013) and MMS (p = 0.017), whereas significant differences for maximum selection coefficient when comparing galactose to MMS (p = 0.008) and DMSO (p = 0.027), and Benomyl to MMS (p = 0.027). The *ESP1* fitness function becomes flatter in some conditions but the deleteriousness of high expression remains important in each case. Therefore, a duplication event could be more or less beneficial depending on the environment, affecting its probability of fixation. The differences between the shape of the fitness function of *ESP1* across environments could explain why its duplication is not observed in yeast strains, even though it would be beneficial in some conditions.

## Discussion

While several examples of positive selection on gene duplication have been proposed to be related to dosage effects (Hardwick et al., 2014; Perry et al., 2007), very few studies have examined how this increase leads to higher fitness relative to what optimal expression would be. In principle, gene duplication leads to a doubling of gene dosage. As long as the fitness of the resulting change in expression is higher than the single-copy expression level, the duplication will be favoured by natural selection. Protein abundance resulting from doubling gene dosage does not need to be at the exact optimal dosage for this to happen; it could be below this value or even overshoot the optimum (Cisneros, Dibyachintan, et al., 2026). Here, we used a graded series of synthetic promoters to characterize the fitness function of genes reported to be beneficial when duplicated in *S. cerevisiae* and examine these possibilities.

We examined a set of 11 genes that were previously shown to improve competitive growth when duplicated on a centromeric plasmid in yeast (Ascencio et al., 2021). Unlike what was found in previous studies (Ascencio et al., 2021; Keren et al., 2016), the overexpression of most of the assayed genes led to little to no effect on fitness. There are many reasons that could explain this result, the most important being that the extent of expression increase was insufficient to alter fitness for several assayed genes. Protein abundance measurements revealed that expression increase varies between genes and that the strongest synthetic promoters often failed to increase protein abundance beyond what is seen when the gene is duplicated with its own promoter. This suggests that other synthetic plasmids may be necessary to achieve higher expression for some of these genes, and that expression is regulated post-transcriptionally to different levels depending on the gene. Additionally, the plasmid system with synthetic promoters presents several differences with a situation where the entire gene would be duplicated (Ascencio et al., 2021), such as the lack of the native untranslated regions in 5’ (5’UTR) and 3’ (3’UTR), which play an essential role in post-transcriptional regulation through effects on translation initiation, localization of mRNA, and mRNA degradation (Leppek et al., 2018; Mayr, 2019). Terminators have indeed been shown to impact mRNA half-life and protein abundance, this effect being especially important when coupled with low-activity promoters (Curran et al., 2013; Yamanishi et al., 2011, 2013). As our plasmid system used the same terminator for each gene instead of the native one, our system could overlook part of the native regulation mechanisms and impact the expression reached at duplication in this study.

One gene previously shown to confer a fitness advantage when duplicated, but that was found to reduce fitness in our experiments, is *GCR1*. Although expression toxicity was observed in most conditions tested, adding agitation to the experimental conditions resulted in entirely flat fitness function. While we did not explore the effect of this condition further, this difference is likely linked with limited media aeration, as the toxicity of overexpression was also not observed in high-throughput studies performed in smaller media volume (Ascencio et al., 2021), with agitation (Keren et al., 2016), or solid media (Arita et al., 2021), possibly due to higher oxygen access. Since toxicity was only observed for *GCR1*, this effect of overexpression is likely to arise from the interaction between limited agitation and gene function, presumably through aeration. Without pointing at the exact cause of the discrepancy between studies, this result shows that different experimental designs can significantly change the relationship between fitness and expression levels for some genes. This, plus the fact that our construct here only includes the coding sequence of *GCR1*, contrary to Ascencio et al. (2021) who duplicated the entire gene region upstream and downstream, could explain the differences between our two studies.

For *ESP1*, we found that increasing expression over the native level can indeed increase fitness and that its duplication leads to an expression level that is slightly higher than the optimum. *ESP1* encodes a separase, which plays an important role in the cell cycle by cleaving cohesin, resulting in the separation of sister chromatids at the initiation of anaphase (Baum et al., 1988; Uhlmann et al., 1999). Esp1p also has a role in the release of Cdc14, a phosphatase contributing to mitotic exit (Sullivan & Uhlmann, 2003). We therefore hypothesized that increased expression of *ESP1* leads to an acceleration of the cell cycle, leading to a proliferation advantage. This was supported by the negative relationship that we found between expression and cell size. Along with the lower proportion of time spent in G2 observed when its expression is increased, this indicates that the overexpression of *ESP1* could result in a faster transition from G2 to G1, but also smaller cell size, providing a mechanistic explanation for the fitness increase through gene duplication of *ESP1*. This could also explain why fitness eventually decreases at high expression levels, if the accelerated cell cycle leads to excessively small cell size and bypasses control checkpoints that ensure appropriate cell size or other important molecular phenotypes before division. The presence of an evolutionary tradeoff related to *ESP1* overexpression is further supported by a previous report of *ESP1* overexpression leading to cell cycle defects such as the premature cohesin cleavage, incorrect spindle assembly, as well as a bypass of the DNA damage checkpoint (Tinker-Kulberg & Morgan, 1999). Over short periods of time such as in a fitness assay, it is therefore possible that these deleterious effects are counterbalanced by the proliferation advantage related to cell division, but that cellular defects related to cell size reduction, aneuploidies or DNA damage could accumulate overtime, resulting in purifying selection on the duplicate that would favor a return to a single copy. Future directions regarding the study of the duplication of *ESP1* effects could therefore focus on fitness effects over longer periods of time to elucidate if the observed fitness advantage persists over time.

For *PDC2*, we also observed increased fitness with overexpression, with the expression at the duplicated state lower than the potential optimum. While we didn’t investigate this case further because of the insufficient overexpression range to observe its optimum, this result provides an interesting contrast to the results of *ESP1* in regards to the positive dosage model. Indeed, while the duplication of each of these genes would be beneficial, the expression levels in the duplicated genotype differ in their relationship with the optimum, one of them exceeding it, the other not reaching it. If mutations increasing expression were to occur after their duplication, they would be favored by natural selection at the native state for both genes, but would have opposite fitness effects following duplication, being favoured following the duplication of *PDC2* and deleterious for *ESP1*. This suggests that the movement along the fitness function through gene duplication can influence the accessibility of mutations differently across genes.

When considering dosage effects, another factor of importance is environmental conditions. Environment-dependent effects of expression changes on fitness have indeed been reported with many different approaches, such as the addition of NaCl in medium in the context of duplication (Ascencio et al., 2021) or copy number variants (Robinson et al., 2021), changes in the carbon source (Keren et al., 2016; Siddiq et al., 2024), or the addition of drug for genes involved in antifungal resistance (Todd & Selmecki, 2020). For *ESP1*, our results show high similarity for fitness functions across conditions, indicating that movements along the fitness function resulting from a duplication are likely to result in similar effects regarding fitness and the accessibility of mutations increasing or decreasing expression even if environmental conditions change, although the generality of this observation across other genes remains to be tested.

In the context of gene duplication, interpreting fitness functions can often be hard, either because of the impossibility of comparing the fitness function to the expression level resulting from gene duplication or because the full fitness function was not explored, preventing positioning where gene duplication would locate expression level on the fitness function. The availability of a series of synthetic promoters allows the obtention of high-resolution fitness functions, to which the duplicated state can then be compared to study dosage effects in one or multiple conditions. As *ESP1* is the only gene for which its duplication was directly compared to its fitness function, extending this type of investigation to more genes will allow us to examine whether *ESP1* represents a specific case or if it represents a general pattern shared with other genes.

## Methods

### Antibiotics, strains and culture media

Ampicillin (BioShop L, AMP201.25), nourseothricin (Cedarlane Labs, AB-102-25G) and G418 (BioShop, GEN418.10) were solubilized at a 1000X concentration, filtered, and stored at-20 °C.

*Escherichia coli* strain MC1061 (*araD139 Δ(araA-leu)7697 Δ(lac)X74 galK16 galE15(GalS) λ- e14-mcrA0 relA1 rpsL150(strR) spoT1 mcrB1 hsdR2*) was used to perform all clonings. Bacteria were cultivated in 2YT medium (1% yeast extract, 1.6% tryptone, 0.2% glucose, 0.5% NaCl) at 37 °C with agitation at 250 rpm. Solid culture media were prepared in the same manner, adding 2% agar. For bacterial cultures transformed with a plasmid, 100 μg/mL of ampicillin (AMP) was added to the medium.

The *S. cerevisiae* strain R1158 (*URA::CMV-tTA MATa his3-1 leu2-0met15-0*) (Hughes et al., 2000) was used to construct the different strains of this project. The strain was cultivated in synthetic complete (SC) medium buffered to pH = 6 (0.174% yeast nitrogen base without amino acid, 2% glucose, complete drop-out mix, 0.1% monosodium glutamate, 1% succinic acid, 0.6% sodium hydroxide). Solid media were prepared the same way but adding 2% agar. For strains transformed with a plasmid, 100 μg/mL of nourseothricin (NAT) was added to the medium.

### Plasmid backbone and clonings

The plasmid yeast_DualReporter (de Boer et al., 2020) was ordered from Addgene (plasmid #127546) and used as the backbone for the different plasmid constructions. Inserts were amplified with at least 20 bp of homology to the backbone in 5’ and 3’, and both backbone and inserts were purified on magnetic beads (Axygen Axy Prep Mag PCR Clean-Up Kit) and eluted in Tris (20 mM, pH = 6). Inserts were cloned in the corresponding plasmid backbone by Gibson DNA assembly (Gibson et al., 2009) and the resulting plasmids were then obtained by transforming them into competent MC1061 *E. coli* cells prepared using a standard calcium chloride protocol and plated on 2YT + AMP medium. Colonies were retrieved by adding 5mL of liquid 2YT medium to the petri dish and scraping all colonies for transformation of pools, or by retrieving a single colony for unique plasmids. Plasmids were extracted from 5mL liquid cultures grown overnight with an alkaline lysis kit (Geneaid Presto™ Mini Plasmid Kit) and transformed into yeast competent cells (Hughes et al., 2000) using a standard lithium acetate transformation protocol (Gietz & Schiestl, 2007).

### Design and transformation of synthetic promoter pools

The competition and control promoter pools were ordered from IDT. Each pool contained 100 promoters of 223 bp from de Boer et al. (2020). These promoters are composed of an 80 bp sequence (Supplementary File S1) flanked by a distal sequence of 54 bp in 5’ and by a proximal sequence of 89 bp in 3’. For both pools, the promoters were chosen from the high-confidence (at least 100 reads) expression dataset of de Boer et al. (2020). We filtered out promoters containing homopolymers of a size higher than or equal to 6 from the dataset before dividing it in 100 intervals of log10 equal expression range. We randomly chose one promoter from each interval to obtain a pool covering the whole dataset’s expression range. For all promoters of the pools, pairwise identity % never exceeds 50%.

Promoter pools were diluted at 0.05 ng/μL in H_2_O and amplified for the first time by PCR without the forward oligo. PCR products were then purified on magnetic beads (Axygen Axy Prep Mag PCR Clean-Up Kit) and eluted in 15 μL of Tris (20 mM, pH=6), and a second PCR was then performed using 3 μL of the purified product as well as both forward and reverse oligos. Both amplifications were performed with the following PCR cycles: 3 min at 98 °C, (30 sec at 98 °C, 15 sec at 60 °C, 10 sec at 72 °C) x 5 cycles, 20 sec at 72 °C.

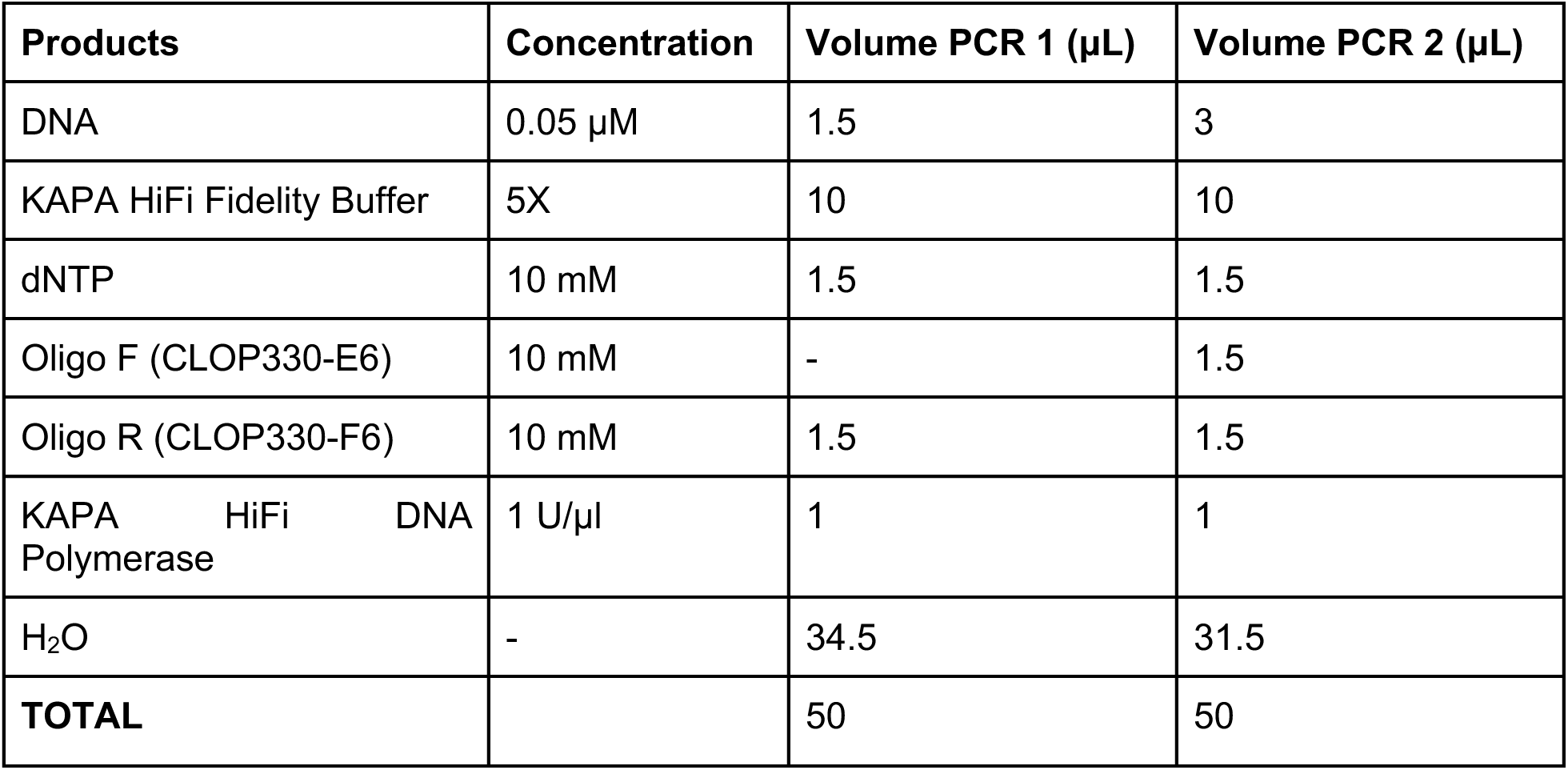

| Products | Concentration | Volume PCR 1 (μL) | Volume PCR 2 (μL) |
| --- | --- | --- | --- |
| DNA | 0.05 μM | 1.5 | 3 |
| KAPA HiFi Fidelity Buffer | 5X | 10 | 10 |
| dNTP | 10 mM | 1.5 | 1.5 |
| Oligo F (CLOP330-E6) | 10 mM | - | 1.5 |
| Oligo R (CLOP330-F6) | 10 mM | 1.5 | 1.5 |
| KAPA HiFi DNA Polymerase | 1 U/μl | 1 | 1 |
| H<sub>2</sub>O | - | 34.5 | 31.5 |
| **TOTAL** |  | 50 | 50 |

The synthetic promoter pools were cloned in the pTpA region of the plasmid and digested with the XhoI restriction enzyme.

### Promoters strength measurements

The gene sequence fragment of the 11 genes studied and their native promoters, defined here as the 1000 bp upstream of the gene start codon in the reference genome, were amplified separately from genomic DNA extracted from the BY4741 *S. cerevisiae* strain. The plasmid was digested with the restriction enzymes BamHI-HF and BmtI-HF, before cloning the native promoters to replace the pTpA region of the yeast_DualReporter, upstream of the yeGFP. Eleven synthetic promoters from the competition pool were ordered as gblocks gene fragments (IDT) and cloned upstream of the yeGFP of the original yeast_DualReporter plasmid digested with the XhoI restriction enzyme. The cloning of these synthetic promoters was validated by Sanger sequencing with the CLOP322-G6 primer.

The expression levels associated with native and synthetic promoters were measured by flow cytometry. Plasmids carrying the promoters upstream of the yeGFP were transformed in the R1158 yeast strain and 8 precultures were incubated overnight at 30 °C in SC + NAT for each transformation. Cultures were diluted in fresh media to 0.15 OD and grown until they reached 0.5 OD. These cultures were diluted 1:10 in sterile H_2_O before measuring the yeGFP’s fluorescence (channel peak: 525 nm) through flow cytometry (Guava EasyCyte HT). For each sample, a minimum of 2000 events were acquired. As a quality control, events with mCherry fluorescence measures (channel peak: 620 nm) lower than 4 units were removed from the analysis (Fig. S7A). Expression for every event was then calculated as the yeGFP fluorescence (channel peak: 525 nm) divided by the forward scatter (FSC) to correct for cell size. The expression measure for each replicate (n=8) of each genotype was calculated as the median of all events.

### Competition assay control and experimental plasmids

The control plasmid used in competition assays, pKB134, was constructed by removing the yeGFP of the yeast_DualReporter, in which synthetic promoters could be cloned without a downstream gene or fluorescent protein. We replaced the yeGFP with the FseI and XbaI restriction enzyme sites to facilitate further plasmid constructions. To do so, the pTpA region and part of the NAT marker were both amplified from the yeast_DualReporter plasmid using the primers CLOP322-G6/CLOP346-G8 and CLOP346-F8/CLO01-37 respectively. The two amplified fragments were then mixed and diluted 1:40 in H_2_O before performing a PCR fusion with the primers CLOP322-G6 and CLO01-37, digesting the fused product with DpnI. The yeast_DualReporter was digested with the BmtI-HF and BglII restriction enzymes, and the digested plasmid was migrated on a 1% agarose gel to extract the fragment corresponding to the backbone without the yeGFP by gel extraction (NEB Monarch Gel extraction Kit). This backbone and the fused product were then assembled to form the control plasmid backbone. The modified fragment of the plasmid was validated by Sanger sequencing with the primer CLOP346-H8. The control pool of plasmids was obtained by digesting this backbone with XhoI and by cloning the control pool of 100 synthetic promoters in the pTpA.

For the construction of the plasmids combining the genes and the synthetic promoters, each gene’s sequence was amplified from the genomic DNA of BY4741 and cloned downstream of the pTpA in the pKB134 plasmid, previously digested with the XbaI restriction enzyme. The competition pool of 100 synthetic promoters was then cloned in the pTpA of the resulting plasmid digested with the XhoI restriction enzyme. For each library described in this section, plasmids extracted from two colonies were validated by whole plasmid sequencing (FlowGenomics).

### Growth curves of yeast in different conditions

Three precultures of the R1158 strain carrying the empty plasmid (pKB134) were grown overnight in SC + NAT media. The following day, 800 μL of media was prepared for 4 concentrations of MMS and 6 concentrations of Benomyl by mixing 792 μL of SC + NAT to 8 μL of the corresponding chemical diluted in DMSO to the concentration needed to obtain 1.11X of the desired media final concentration. 180 μL of media was then transferred in triplicate in a deep-well 96 plate. The OD of the precultures was then measured and cells were then diluted so that 20 μL could be added to the plates to obtain an OD of 0.1. The plate was then incubated without agitation at 30°C in a Epoch2 plate reader (Agilent-BioTek) for 48 hours, measuring OD every 15 minutes. From this data, growth curves were obtained by fitting a generalized additive model for each condition. The concentration used for the competition assay for both MMS and benoyml were chosen by comparing the growth curves of each concentration visually and by calculating the area under the curve (AUC), aiming for a slight decrease in fitness relative to SC + NAT + DMSO (Fig. S8), AUC being calculated using the trapezoidal rule as followed:

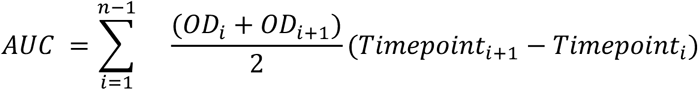

### Competition assays

For each of the 11 genes selected, the experimental plasmid pool was mixed with the control plasmid pool in a 1:1 ratio, resulting in a pool in which the 200 different plasmids are in equal concentration. The mixed plasmid pools were then transformed into competent cells of the standard R1158 *S. cerevisiae* strain and aimed to obtain at least 2000 colonies. The transformed colonies were mixed, and three independent liquid cultures were started in 5 mL of SC + NAT medium to grow overnight. The next day, a 1:25 dilutions of these cultures were prepared in 5 mL of fresh media, which we identified as time point 0 for the competition assay. We let these cultures grow for 24 hours at 30 °C without agitation before redoing new 1:25 dilutions daily for 7 days, which corresponds to about 27 generations. For each time point, 2 samples of 5 OD were taken for each culture, and the cells were kept at-80 °C until further manipulations.

For *ESP1* and *GCR1*, assays were also performed using the same libraries of plasmids, but with the following modifications to the experimental conditions: galactose: (2% galactose instead of glucose), 50% glucose (50% reduction of all media components), raffinose (2% raffinose instead of glucose), agitation (SC + agitation), benomyl (SC + 5 μg/mL benomyl), methyl methanesulfonate (MMS) (SC + 0.001% MMS) and DMSO (addition of 80 μL of DMSO to the media), the last one being the control for benomyl and MMS.

### Libraries preparation

For each sample, 5 OD units were transferred in a 1.5 mL tube and centrifuged at 600 g for 2 minutes to isolate cell pellets. These pellets were then stored at-80 °C until further manipulations. Plasmid extractions were performed using Zymoprep Yeast Plasmid Miniprep II kits, with modifications following recommendations from previous studies (Bédard et al., 2024; Starr et al., 2020). Pellets were resuspended in 20 μL of Solution 1 and 30 U of zymolyase™ was then added before incubating for 2 hours at 37 °C. Tubes were then stored again at-80 °C overnight before performing the next steps. Samples were thawed at 37 °C for 3 minutes before adding 200 μL of Solution 2 and 400 μL of Solution 3, mixing well after each addition. Samples were centrifuged at maximal speed for 3 minutes, and the supernatant was then transferred in a new 1.5 mL tube before centrifuging again for 3 minutes at maximal speed. Supernatant was transferred to Zymo-Spin-I columns before a 30-second centrifugation at maximal speed. 550 μl of Wash Buffer was then added to the columns before centrifuging at maximal speed for 30 seconds. Columns were dried by centrifuging them again for 90 seconds at maximal speed. The content of the column was finally eluted in 10 μL of Tris (10mM, pH = 8), and the eluant was conserved in 1.5 mL tubes at-20 °C.

The synthetic promoters in the plasmids were then amplified using 5 μL of each miniprep and the following PCR cycles: 3min at 98°C, (30 sec at 98°C, 15 sec at 61°C, 20 sec at 72 sec) x 18 cycles, 30 sec at 72°C.

| Products | Concentration | Volume (µL) |
| --- | --- | --- |
| Miniprep | 1 ng/µl | 5 |
| KAPA HiFi Fidelity Buffer | 5X | 5 |
| dNTP | 10 mM | 0.75 |
| Oligo F (CLOP333-C2) | 10 mM | 0.75 |
| Oligo R (CLO6-64) | 10 mM | 0.75 |
| KAPA HiFi DNA Polymerase | 1 U/μl | 0.5 |
| H<sub>2</sub>O |  | 12.25 |
| **TOTAL** |  | 25 |

PCR products were purified on magnetic beads (Axygen Axy Prep Mag PCR Clean-Up Kit) and eluted in 20 μL of Tris (20 mM, pH = 8), and their DNA concentration was measured using a Nanodrop spectrophotometer (Thermo Fisher Scientific). The products were diluted to 0.01 ng/μL in H_2_O and a second PCR was performed to add Illumina indexes with the following PCR cycles: 3min at 98°C, (30 sec at 98°C, 15 sec at 61°C, 20 sec at 72 sec) x 16 cycles, 30 sec at 72°C.

| Products | Concentration | Volume (μL) |
| --- | --- | --- |
| DNA | 0.01 ng/μl | 1 |
| KAPA HiFi Fidelity Buffer | 5X | 5 |
| dNTP | 10 mM | 0.75 |
| Oligo F | 10 mM | 0.75 |
| Oligo R | 5 mM | 1.50 |
| KAPA HiFi DNA Polymerase | 1 U/μl | 0.3 |
| H<sub>2</sub>O |  | 15.70 |
| **TOTAL** |  | 25 |

The PCR products were purified with magnetic beads and eluted in 20 μl of Tris (20 mM, pH = 8), and 5 μL of each sample was then pooled in a 1.5 mL tube before sending it to the sequencing platform of Laval University for Aviti sequencing or to the CHUL sequencing platform for Illumina NovaSeq6000 150 paired-ends sequencing, aiming for 1000 reads per variant.

### Read processing and selection coefficient

From the deep sequencing data, reads which couldn’t be aligned perfectly to one of the promoter sequences of the experiments were excluded and only the promoters presenting at least 100 reads at the first time point (T0) were processed to avoid noisy measurements due to low read depth. From the read counts, we calculated the proportion of reads associated with each promoter at a given time point. For each promoter within each culture, a selection coefficient was calculated using the following formula:

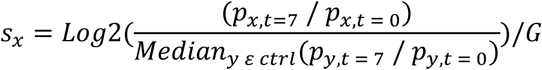

Where p_x, t_ is the proportion of reads associated to a promoter_x_ at given time point (t), p_y, t_ is the proportion of reads associated to a promoter_y_ of the control (ctrl) pool at a given time point, and *G* is the generation time between the two time points.

### Fitness functions

The selection coefficient calculated for each strain was associated with the reported expression measures (de Boer et al., 2020) to obtain the fitness function of each gene for the expression range conferred by the plasmid gene copy. A LOESS regression was fitted (frac = 0.4) using the lowess function from the Python package statsmodels (version 0.14.0) (Seabold & Perktold, 2010) and the 95% confidence interval of the function was calculated using a bootstrap (n = 200) approach. As all fitness functions presented a bias between the selection coefficient values of the control and experimental plasmids, the initial bias of the curve was calculated as the value of the selection coefficient for the expression of the first promoter. For the assays made in different conditions for the same gene, the curve was adjusted by subtracting this effect from the computed regression values to allow the comparisons of the function in different conditions.

### Protein abundance measurements

To study the protein abundance of different strains, we tagged both the genomic and plasmid gene copies with the same yeGFP from the yeast_DualReporter. To tag the genomic copy of each gene, we amplified the yeGFP from the yeast_DualReporter with 40 bp of homology to the 3’ end of the gene of interest, which we fused with the *HIS3MX6* marker amplified from the pYM31 plasmid (Janke et al., 2004) with 40 bp of homology to the downstream sequence of the gene of interest in 3’ using fusion PCR. The resulting cassettes were then individually inserted into the genome of competent R1158 cells (Hughes et al., 2000) using a standard lithium acetate transformation protocol(Gietz & Schiestl, 2007).

To study the duplicated gene state, we amplified both the native promoter (1000 bp upstream of gene sequence) and gene sequence from the BY4741 *S. cerevisiae* genomic DNA and cloned it in the yeast_DualReporter plasmid backbone, digested with the restriction enzymes BamHI-HF and BmtI-HF to replace the pTpA and fuse the gene with the plasmid’s yeGFP. For the protein abundance of strains with specific synthetic promoters, we amplified the genes of interest and fused them to an amplified the fragment of the yeast_DualReporter containing the yeGFP and ∼650 bp in 3’ of its coding sequence using fusion PCR, which we then cloned in the pKB134 plasmid, digested with the XbaI restriction enzyme. This strategy ultimately resulted in the fusion of the gene sequence with the yeGFP in the plasmid, downstream of the pTpA. The fusion of the gene and the yeGFP was validated by Sanger sequencing with the primer CLOP81-F3. The resulting plasmids could then be digested with the XhoI restriction enzyme to clone the synthetic promoter of interest in the pTpA, validated by Sanger sequencing using the primer CLOP322-G6. The constructed plasmids were transformed in the yeGFP-tagged yeast strains to measure the total abundance of the proteins by flow cytometry using the same method as previously described in the “Native promoter strength measures” section.

For protein abundance levels of *ESP1* with different promoters, two controls were used. One was the *ESP1*::yeGFP R1158 strain transformed with the pFB110 plasmid, which contains a copy of *ESP1* with the first three methionines substituted by stop codons fused to the yeGFP after the empty pTpA to represent the single copy state. The second is the untagged R1158 strain transformed with the pFB109 plasmid, which contains the same *ESP1* sequence as pFB110, but without the yeGFP fusion, and was used as a negative control. These two controls were chosen to avoid biases arising from differences in plasmid size. To construct these plasmids, the 90 first nucleotides of the coding sequence of *ESP1*, with the three first methionines replaced by stop codons, was ordered as a gblock with 35 nucleotides of homology to the pTpA. Starting immediately after the third methionine, the remaining sequence of *ESP1* was amplified either with 5’ homology to the sequence after the pTpA in the pKB134 plasmid (pFB109, oligos CLOP376-H1 and CLOP332-H10) or with the yeGFP (pFB110, oligos CLOP376-H1 and CLOP376-D1). For the pFB110 construction, the fragment of *ESP1* was fused to the yeGFP as described above. The final plasmids were obtained by Gibson assembly using the pKB134 backbone digested with XbaI, the gblock and the remaining *ESP1* sequence fragment. Both plasmids were validated by whole plasmid sequencing.

### Cell size measurements

Size measurement associated with the expression levels of *ESP1* of different synthetic promoters was performed in the R1158 yeast strain transformed with the plasmids containing a gene copy downstream of the pTpA and in which the yeGFP was removed, described in the “Competition assay control and experimental plasmids” section, transformed with synthetic promoters and validated by Sanger sequencing with the primer CLOP322-G6. For the duplicated state, the native promoter and gene of interest were amplified from genomic DNA and cloned in the pKB134 plasmid, digested with the restriction enzymes BmtI and XbaI to replace the pTpA. For the single-copy state, the pFB110 plasmid (described in the previous section) was transformed in the same yeast strain to avoid potential bias resulting from plasmid size. Cell size measurements were obtained by flow cytometry (Guava EasyCyte HT) with a minimum of 5000 events per sample, using the forward scatter channel (FSC) signal as an indicator of cell size. The median was calculated for each replicate for further statistical analysis.

### DNA content measurements and cell cycle estimation

DNA content was measured from fixed cell cultures by adapting the method used in (Bautista et al., 2024). From overnight precultures of the R1158 strain carrying a plasmid containing an extra copy of *ESP1* controlled by a single promoter (8 replicates / genotype), cultures were started at 0.15 OD in 2 mL of SC complete pH = 6 + NAT with agitation at 30°C and grown until they reached 0.5 OD. From each culture, 1.5 mL was spun down (3min, 3000 rpm) and supernatant was removed before fixing the cells in 1 mL of ethanol 70%, keeping cells at 4°C until further manipulation. 25 μl of fixed cells were then harvested (3 minutes, 3000 rpm, removing the supernatant), and washed with 200 μl of sodium citrate (50 mM, pH = 7). Cells were harvested again, and washed two times with 200 μl of sodium citrate + EDTA (30 mM), leaving the resuspended cells at room temperature for 5 minutes after each addition. Cells were harvested again and resuspended in 200 μl of sodium citrate + RNAse (0.25mg/mL) and incubated overnight at 37°C. The next day, cells were harvested and washed two times with 200 μl sodium citrate. In a transparent plate, 100 μl was transferred and mixed with 150 μl of sodium citrate + 1μM SytoxGreen (Invitrogen, #S7020) and kept in the dark at room temperature for 1h before measuring fluorescence in flow cytometry (channel peak: 525 nm), acquiring 10 000 events per sample. Events were gated using the forward and side Scatters (FSC and SSC) as quality control (Fig S7B).

To estimate the proportion of cells in G1 and G2, a gaussian mixture model with the function “normalmixEM” of the mixtools R package (v.2.0.0.1; (Benaglia et al., 2009)) was fitted over the data, using k = 3 to assign events to either 1C, 2C or background noise, the latter accounting for less than 10% of all events for all cultures. The addition of this third category resulted in more consistent results compared to k = 2, which often led to a single gaussian curve spanning the whole fluorescence range and to which most events were assigned. Initial means (chosen after visualizing histograms of the data) were fixed identically for all samples. For each replicate, the weight of the curves representing 1C and 2C were used to calculate a ratio, then used to perform statistics between the different expression levels.

## Data analysis

All data related to measures obtained through flow cytometry were analysed using R Statistical Software (v.4.3.2; R Core Team 2023). Data related to the competition assays were analysed using Python (v.3.11.14; Python Software Foundation). Figures were assembled using Inkscape (v.1.3; Inkscape Project).

## Data and code availability

The sequences and the expression estimates of the synthetic promoters selected from de Boer et al. (2020) for this project are available in the Supplementary file S1. Oligos and PCR conditions used for the different parts of this project are available in the Supplementary file S2. All scripts and files required to reproduce figures and analyse data are available on Github at https://github.com/Landrylab/Bedard_et_al_2026. Sequencing data are available at the NCBI Sequence Read Archive under BioProject PRJNA1508198: http://www.ncbi.nlm.nih.gov/bioproject/1508198.

## Author contributions

FB, CRL, AKD and IGA designed the study. FB performed the experiments, as well as the data analyses. FB and CRL wrote the manuscript.

## Acknowledgements

We thank S. Aubé and P. Venkataraman, who provided helpful comments on this manuscript. This work was supported by a NSERC discovery grant RGPIN-2026-06411 and a Human Frontiers Science Program grant RGP011/2025 to CRL. CRL holds the CIHR Canadian Research Chair in Cellular Systems and Synthetic Biology. FB was supported by a master’s training fellowship from Fonds de recherche du Québec - Nature et technologies and a Leadership and Commitment scholarship from Université Laval.

## Supplementary Material

**Supplementary Table 1.**
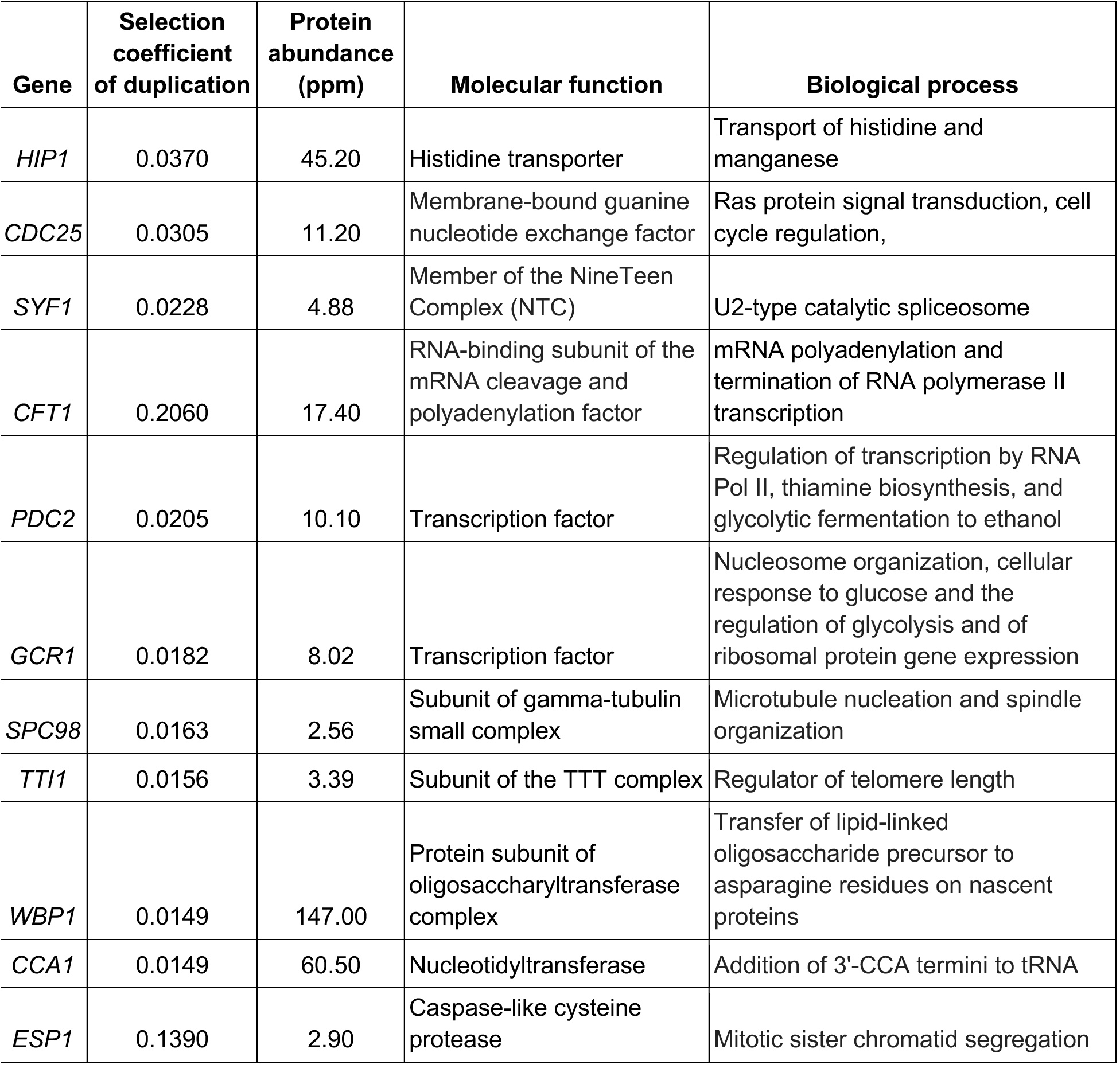
Description of the 11 genes studied. Selection coefficients associated with the duplication of each gene were obtained from Ascencio et al. (2021). Protein abundances were retrieved from integrated abundance estimates available on PaxDb (Huang et al., 2023). Molecular functions and biological processes were obtained from the Saccharomyces Genome Database (SGD).

## List of supplementary tables and figures

**Supplementary Figure 1.**
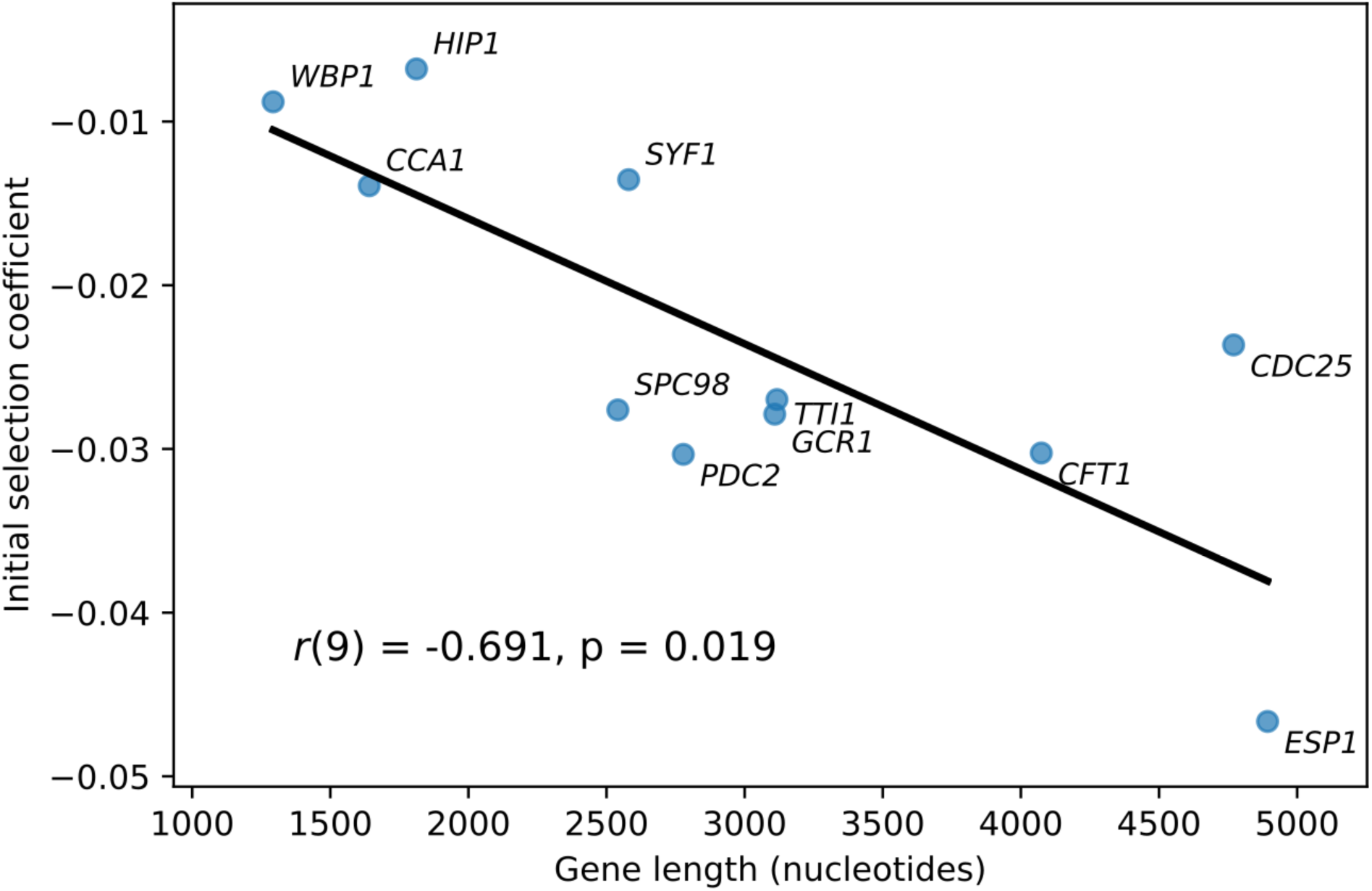
The initial selection coefficient of fitness functions correlates with the length of the gene of interest. The initial selection coefficient, computed from the modelled fitness functions at the promoter strength of the weakest promoter, was calculated for the 11 genes studied. Spearman correlation coefficient and p-value are shown.

**Supplementary Figure 2.**
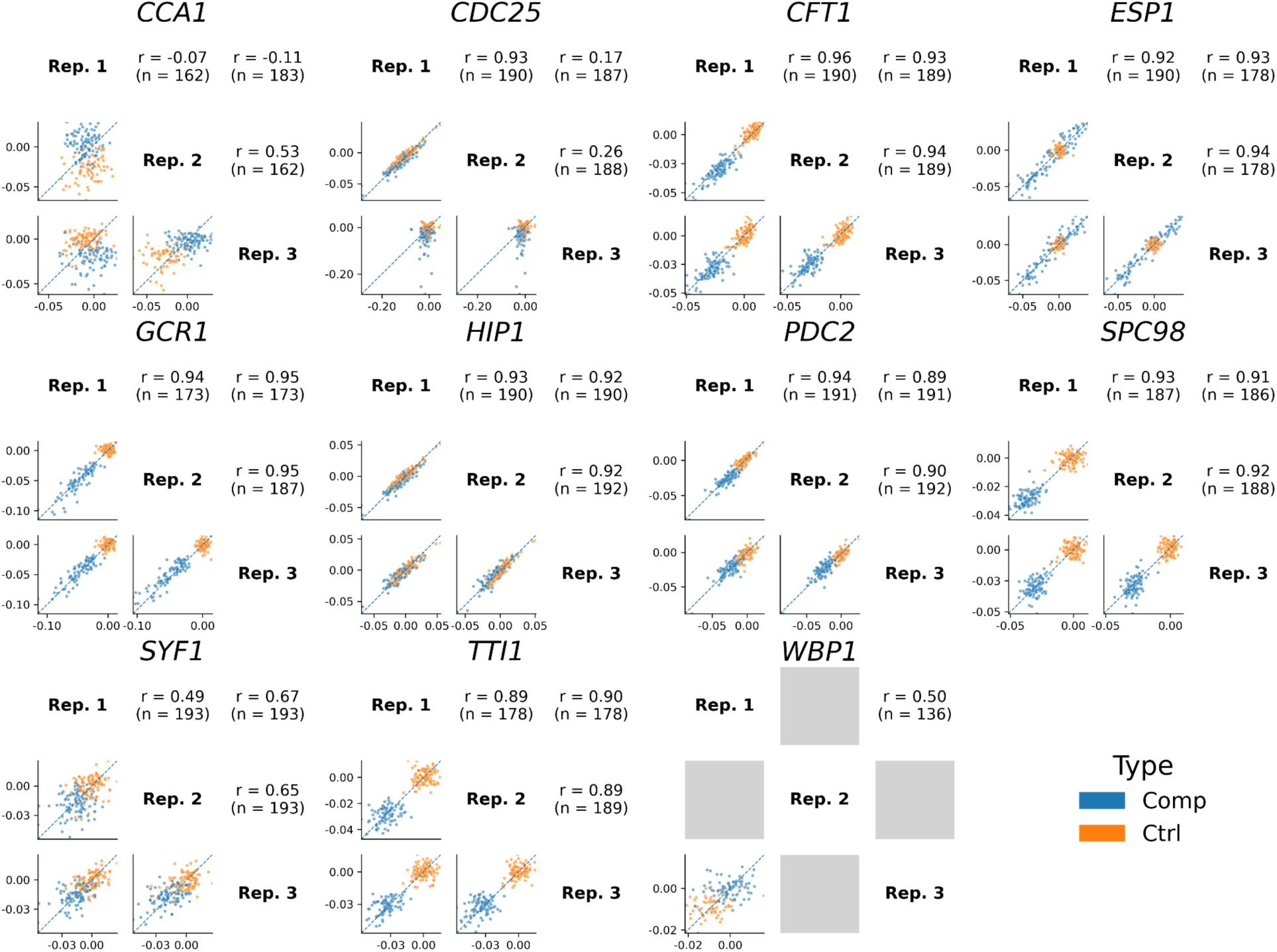
Replicability of selection coefficients between replicate cultures for the competition assays of 11 genes in SC complete media. Pearson correlation between each replicate pair is indicated, as well as the number of promoters for which a selection coefficient was calculated in both replicates (n). Gray squares represent comparisons in which fewer than 100 data points were available.

**Supplementary Figure 3.**
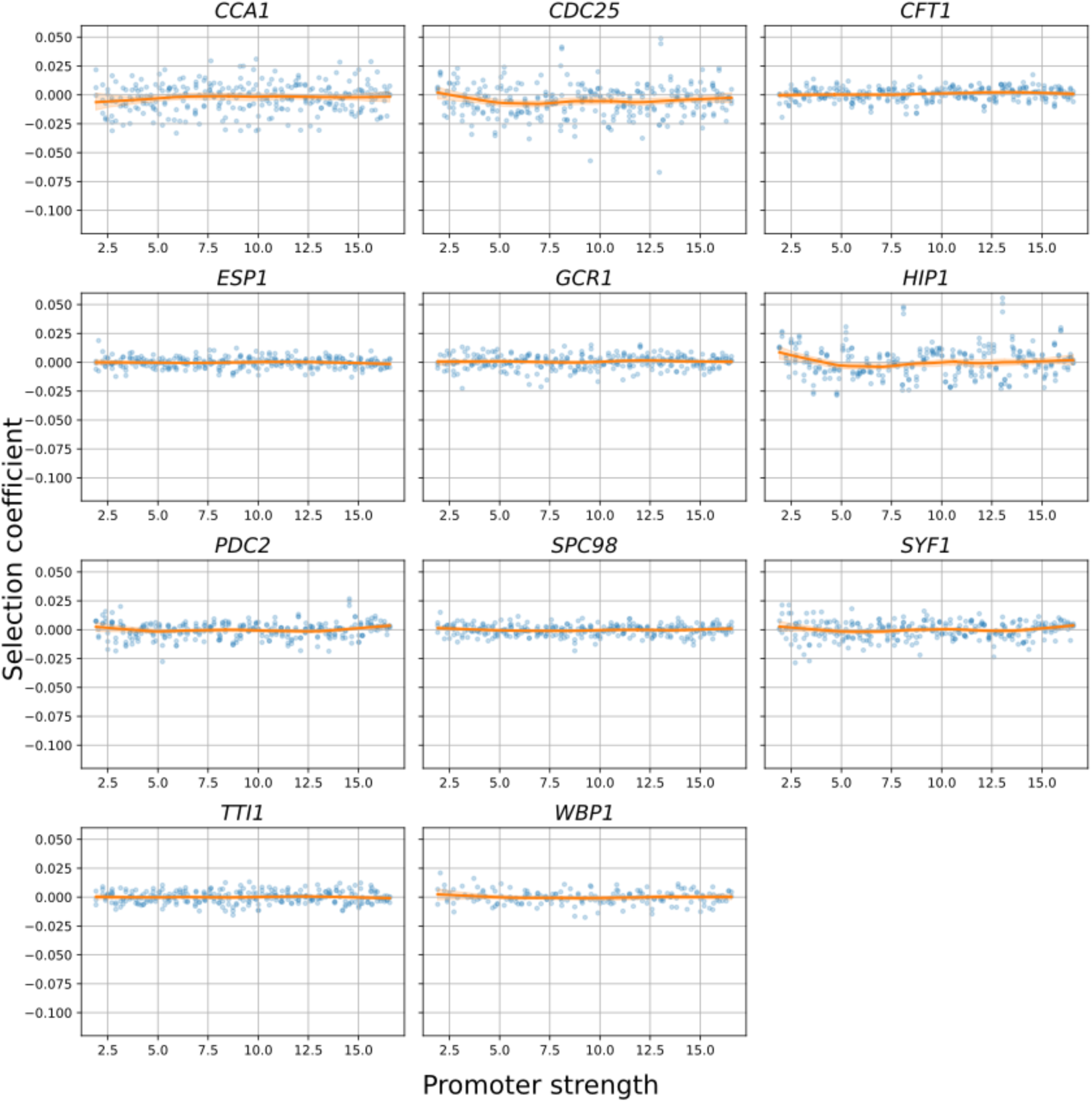
The control plasmids containing the synthetic promoters but no coding sequence exhibit constant selection coefficients for all genes. Fitness functions were obtained using a LOESS regression (frac = 0.4) and shown in orange, whereas the shaded areas represent the bootstrap 95% confidence interval (n = 200). Selection coefficients were calculated for each promoter across all replicates (n = 3) and are shown as faded points.

**Supplementary Figure 4.**
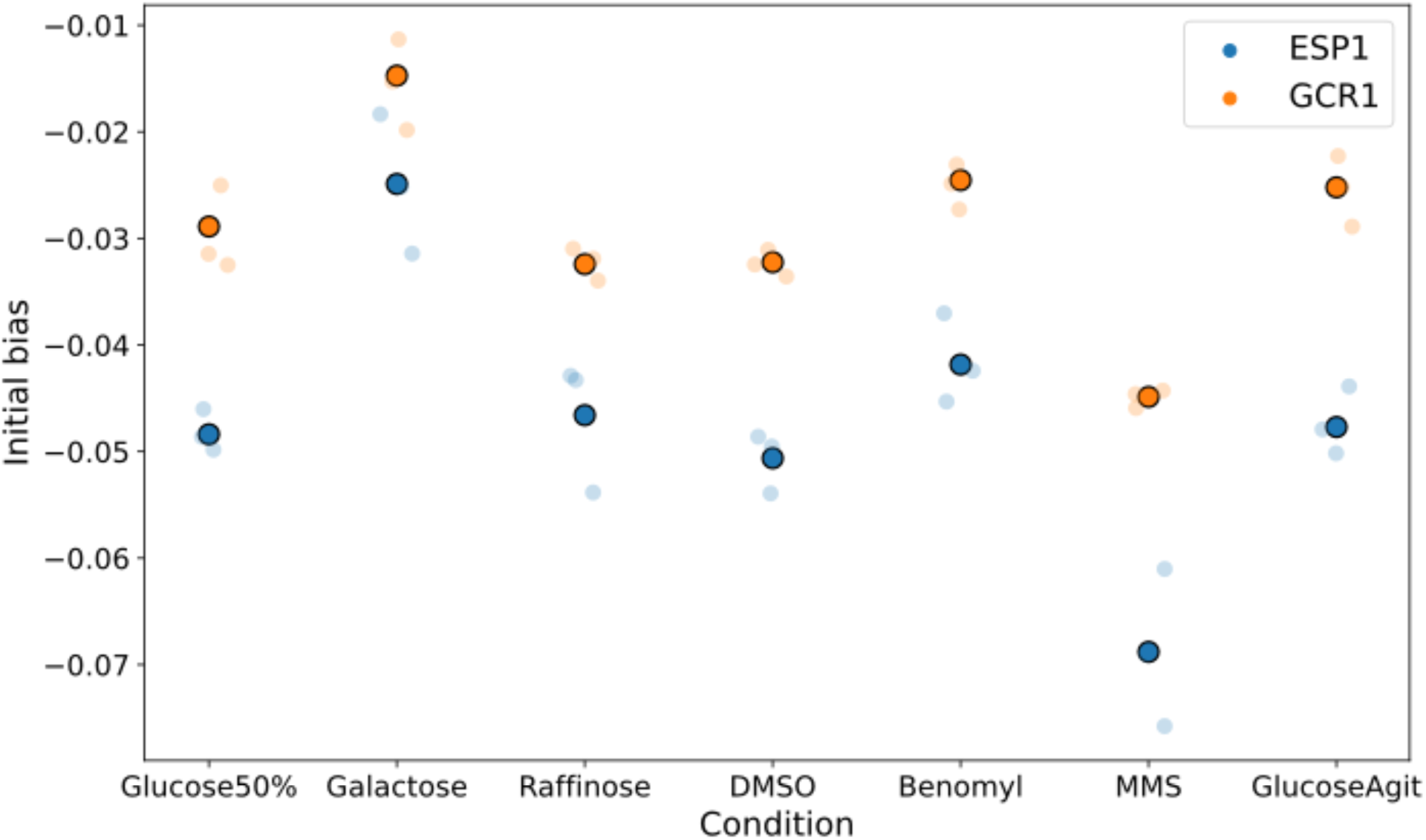
Initial selection coefficient bias across conditions for *ESP1* and *GCR1*. Solid points represent the initial bias calculated across all replicates, whereas faint points represent bias for each replicate (n=3) individually.

**Supplementary Figure 5.**
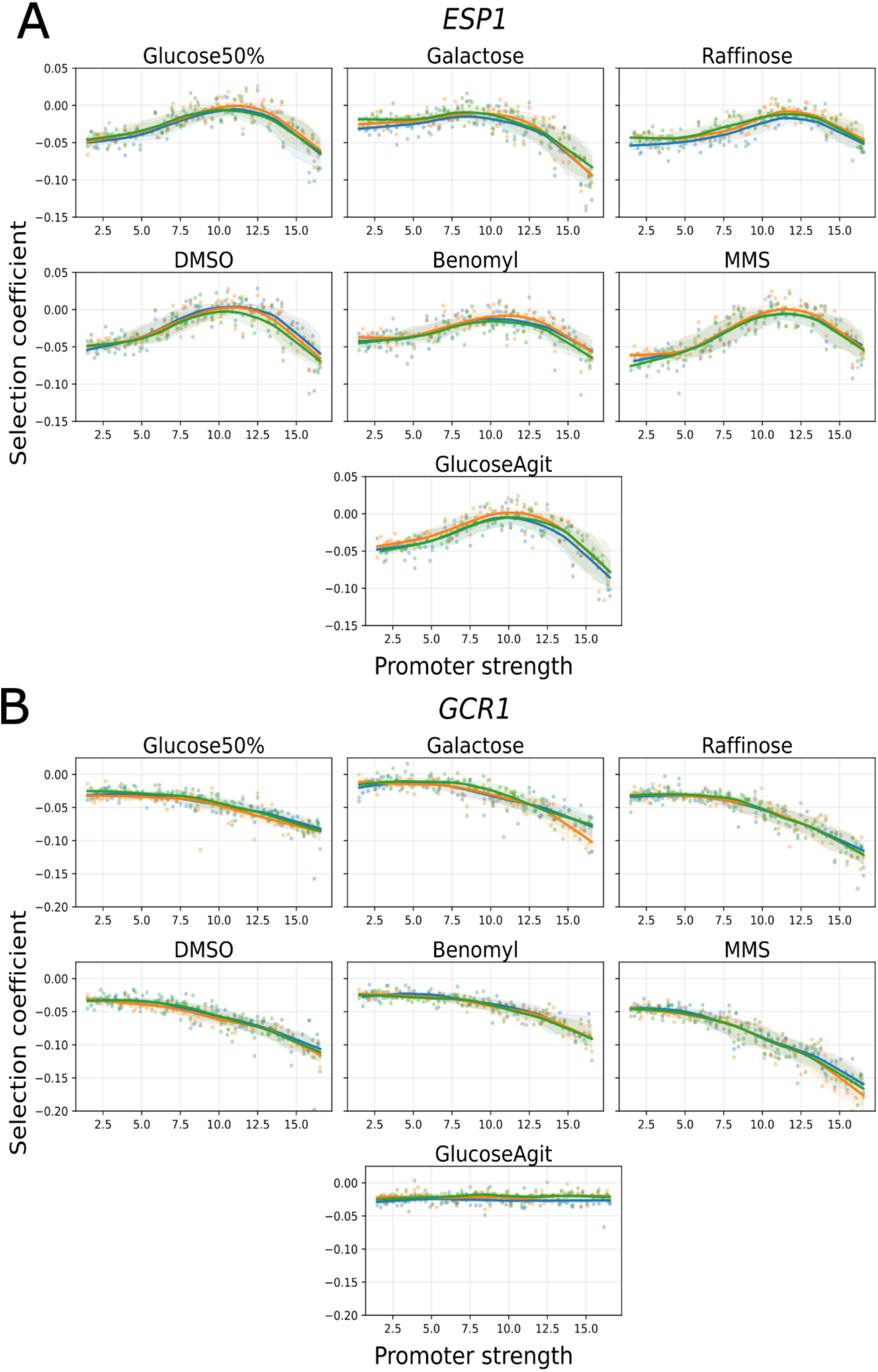
Uncorrected fitness functions of *ESP1* and *GCR1* measured in different experimental conditions for each replicate. Replicates are identified with different colors).

**Supplementary Figure 6.**
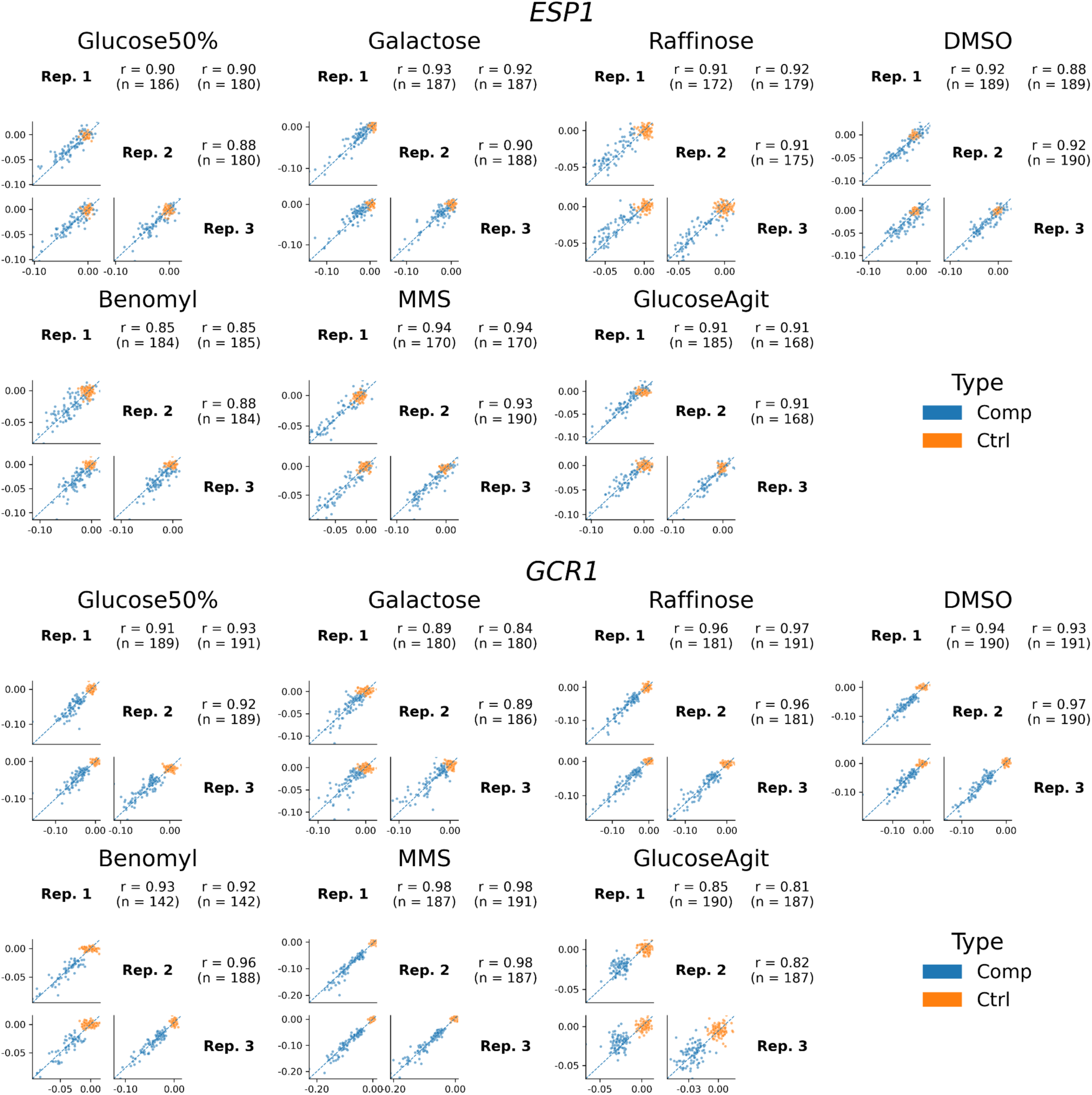
Replicability of selection coefficients between replicate cultures for the competition assays of *ESP1* and *GCR1* in multiple conditions. Pearson correlation between each replicate pair is indicated, as well as the number of promoters for which a selection coefficient was calculated in both replicates (n).

**Supplementary Figure 7.**
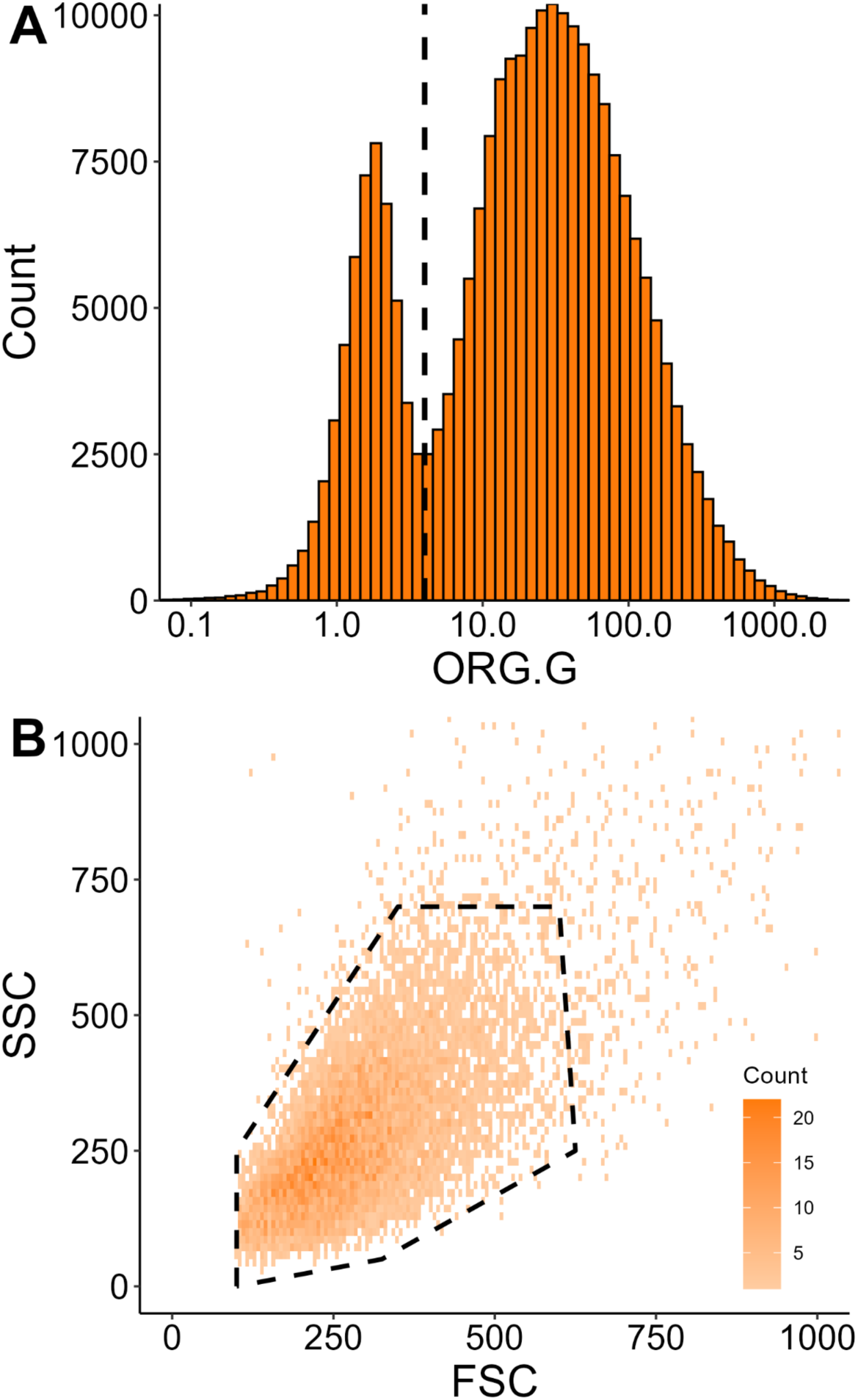
Examples of flow cytometry data gating for A) protein abundance and cell size measurements and B) DNA content measurements. The former aims to remove events with low mCherry fluorescence (ORG.G < 4). This limit is identified as the vertical dashed line on the distribution of fluorescence for the dataset of synthetic promoter strength (presented in Fig. 2A). The latter aims to remove events with cells of unusual size or clusters of cells when measuring DNA content (Fig. 5F), the limits being represented as a dashed polygon on the data of a single sample chosen randomly (pSyn65, rep. 6) for better visualization.

**Supplementary Figure 8.**
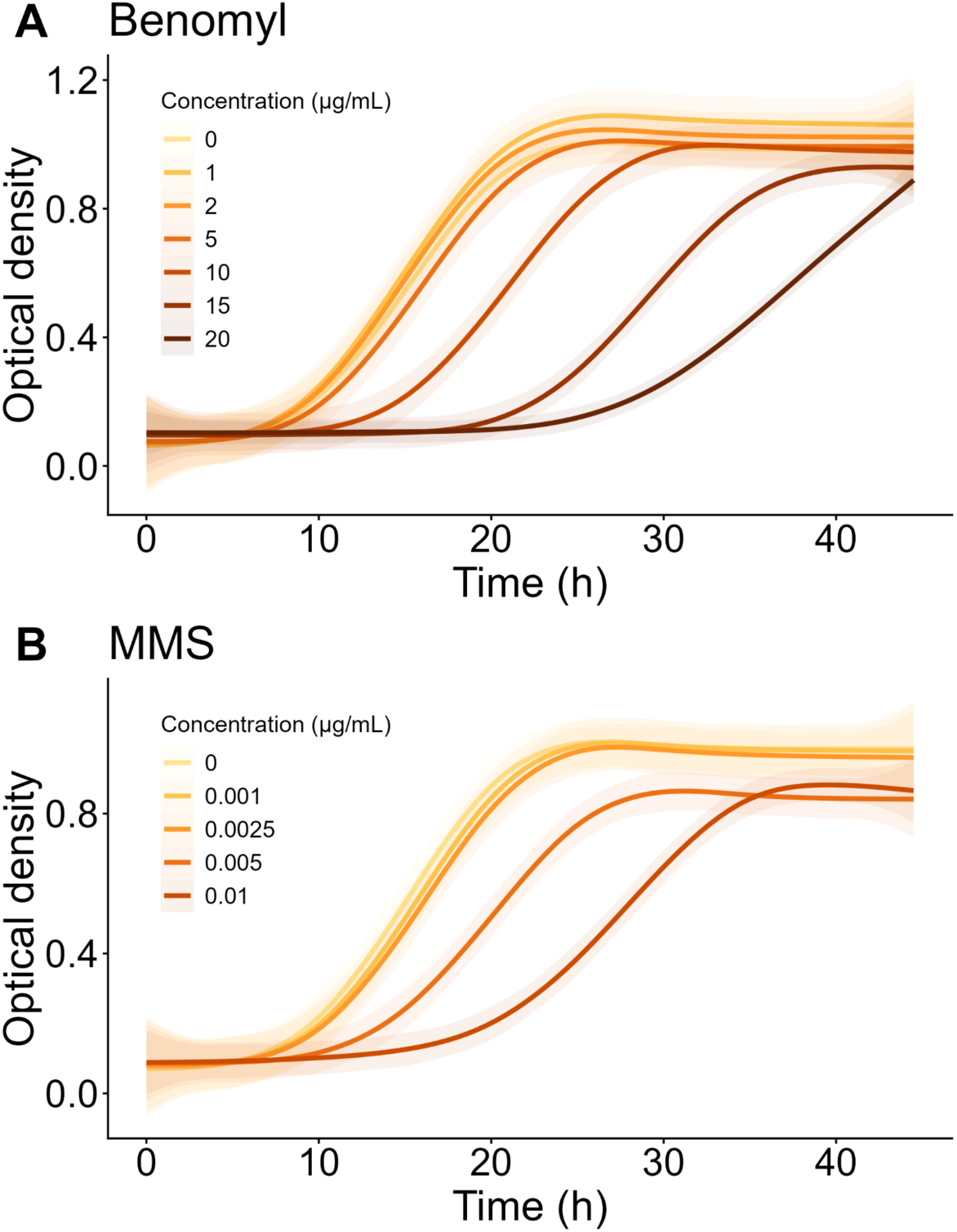
Growth curves of R1158 + pKB134 (empty plasmid) from cultures in SC pH = 6 + NAT and different concentrations (conc.) of A) Benomyl and B) MMS. Three replicates were obtained per condition. Shaded areas around the curves represent the 95% confidence intervals.

